# Downregulation of Stem-loop binding protein by nicotine *via* α7-nicotinic acetylcholine receptor and its role in nicotine-induced cell transformation

**DOI:** 10.1101/2022.03.21.485231

**Authors:** Qi Sun, Danqi Chen, Amna Raja, Gabriele Grunig, Judith Zelikoff, Chunyuan Jin

## Abstract

The use of electronic-cigarettes (e-cigs) has increased substantially in recent years, particularly among the younger generations. Liquid nicotine is the main component of e-cigs. Previous studies have shown that mice exposed to e-cig aerosols developed lung adenocarcinoma and bladder hyperplasia. These findings implicated a potential role for e-cig aerosols and nicotine in cancer development, although the underlying mechanisms are not fully understood. Here we report that exposure to liquid nicotine or nicotine aerosol generated from e-cig induces downregulation of Stem-loop binding protein (SLBP) and polyadenylation of canonical histone mRNAs in human bronchial epithelial cells and in mice lungs. Canonical histone mRNAs typically do not end in a poly(A) tail and the acquisition of such a tail via depletion of SLBP has been shown to causes chromosome instability. We show that nicotine-induced SLBP depletion is reversed by an inhibitor of α7-nAChR (nicotinic acetylcholine receptors) or siRNA specific for α7-nAChR, indicating a nAChR-dependent reduction of SLBP by nicotine. Moreover, not only CDK1 and CK2, two kinases well known for their function for SLBP phosphorylation and degradation, but also CDK2 and PI3K/AKT pathways are shown to be involved, α7-nAChR-dependently, in nicotine-induced SLBP depletion. Importantly, nicotine-induced anchorage-independent cell growth is attenuated by inhibition of α7-nAChR and is rescued by overexpression of SLBP. We propose that the SLBP depletion and polyadenylation of canonical histone mRNAs *via* activation of α7-nAChR and a series of downstream signal transduction pathways, are critical for nicotine-induced cell transformation and potential carcinogenesis.

## Introduction

Electronic cigarettes (e-cigs) are battery-operated devices that deliver nicotine in aerosols by electronic heating of a solution usually containing nicotine, solvents propylene glycol (PG) and vegetable glycerin (VG), and flavorings. Nicotine itself has generally been considered non-carcinogenic until recently. As a consequence, e-cigs are often used as an alternative for conventional cigarettes and thus the use of e-cigs has increased substantially, particularly amongst younger sub-populations (1,2). However, growing numbers of recent studies have recently demonstrated diverse health effects of e-cigs, including their potential carcinogenicity. For example, mice exposed to e-cig aerosols developed lung adenocarcinoma and bladder urothelial hyperplasia (3,4). This implicated a potential role for e-cig aerosols and nicotine in lung cancer development. Specifically, genotoxicity was proposed as the cause of nicotine-induced tumorigenesis based on the observation that e-cig aerosols induce mutagenic DNA adducts, i.e., cyclic 1, N^2^-γ-hydroxy-propano-deoxyguanosine [γ-OH-PdG] and O^6^-methyldG, and inhibits DNA repair in the lungs of ECS-exposed mice and in human lung epithelial cells (3,4).

In addition to mutagenic effects, nicotine is believed to exert tumorigenic effects via nicotinic acetylcholine receptors (nAChRs)-mediated activation of a variety of downstream signaling pathways (5,6). Nicotinic AChRs are expressed in non-neuronal cells, including lung epithelial cells (7). There is a growing body-of-evidence demonstrating that nicotine binds to nAChRs with a higher affinity than acetylcholine and induces overexpression of nAChRs. Moreover, different nAChR subtypes are overexpressed in various cancers and cancer progression is known to be associated with overexpression of nAChRs (6). The importance of nAChRs in cancer and in nicotine-induced carcinogenesis is further supported by the findings that nAChR genetic variants increase the risk of developing cancer, and that some single nucleotide polymorphisms (SNPs) in nAChRs regulate the sensitivity to nicotine (5,8). Major tumorigenic effects of nicotine and principal downstream pathways mediated by nAChRs include, but are not limited to, facilitated cancer cell growth and proliferation by JAK2/STAT pathway, as well as inhibition of apoptosis by PI3K/AKT pathway, regulation of cancer cell detachment, migration and re-attachment by Ca^2+^-dependent recruitment of CAMKII and PKC and subsequent phosphorylation and dephosphorylation of adhesion and cytoskeletal proteins (9–11). While numerous signaling pathways, including those above-mentioned, are implicated in nicotine-induced toxicity and carcinogenicity, the effector(s) that function at ‘final’ steps in the nAChR-mediated signaling pathways have remained elusive.

Previously we have demonstrated that exposure to arsenic, a heavy metal carcinogen, downregulates the expression of Stem-loop binding protein (SLBP), leading to polyadenylation of canonical histone H3.1 mRNA (12). This was surprising since the expression of replication-dependent canonical histone genes, such as histone H3.1, is largely limited in S phase and their mRNAs do not contain a typical poly(A) tail at their 3’ ends like other metazoan genes. Instead, they contain a highly conserved hairpin structure, which is bound by SLBP for proper 3’ mRNA processing, generating mature H3.1 mRNA without a poly(A) tail. Mutations or depletion of this protein can result in misprocessing of the canonical histone mRNAs, leading to expression of polyadenylated mRNA from each of the canonical histone genes (13–15). Unlike normally processed canonical histone mRNAs, polyadenylated mRNAs are relatively stable, which results in existence of canonical histone mRNAs outside of the S phase and increase in H3.1 protein level (16,17). Polyadenylation of H3.1 mRNA has been shown to lead to displacement of histone variant H3.3 at active critical gene regulatory regions, causing deregulation of cancer-related genes, cell-cycle arrest, chromosome instability and cell transformation (17). These data indicate that overexpression of polyadenylated H3.1 mRNA due to SLBP depletion has potential carcinogenic effects (12,17).

In this study, we investigated whether nicotine induces downregulation of SLBP and polyadenylation of canonical histone H3.1 mRNA in human bronchial epithelial cells and in mice, and whether nicotine-induced SLBP depletion is nAChRs dependent. Moreover, we characterized the factor(s) and signaling pathway(s) involved in nicotine-induced downregulation of SLBP. as well as the role of nAChR activation and SLBP depletion in nicotine-induced cell transformation in cultured human lung epithelial cells.

## Materials and Methods

### Reagents

Nicotine (^#^N3876), 2-dimethylamino-4,5,6,7-tetrabromo-1H-benzimidazole (DMAT, ^#^SML2044), and Roscovitine (^#^R7772) were purchased from Sigma-Aldrich (St. Louis, MO). Trizol reagent was obtained from Invitrogen (Thermo Fisher Scientific, Waltham, MA). The enhanced chemiluminescence (ECL) plus kit was purchased from Bio-Rad (Hercules, CA). The bicinchoninic acid (BCA) protein assay kit was obtained from Pierce (Thermo Fisher Scientific, Waltham, MA). The antibodies were purchased as follows: anti-CDK2 (1:1000 dilution, ^#^ab235941), anti-α7-nAChR (1:1000 dilution, ^#^ab216485), anti-histone H3 (1:2000 dilution, ^#^ab1791), and anti-SLBP (1:2000 dilution, ^#^ab181972) were from Abcam (Cambridge, MA); anti-CDK1 (1:1000 dilution, ^#^33-1800) and anti-CK2 (1:1000 dilution, ^#^PA5-95701) were purchased from Invitrogen (Thermo Fisher Scientific, Waltham, MA); anti-histone H4 (1:500 dilution, ^#^05-858), anti-histone 2A (1:500 dilution, ^#^07-146), and anti-histone 2B (1:500 dilution, ^#^07-371) were from MilliporeSigma (Burlington, MA); anti-GAPDH (1:2000 dilution, ^#^sc-47724) was from Santa Cruz Biotechnology (Dallas, TX). Antibodies against phosphoylated-AKT^T308^ (p-AKT^T308^, 1:500 dilution, ^#^2965), p-AKT^S473^ (1:500 dilution, ^#^4060), and AKT (1:1000 dilution, ^#^4961s) were from Cell Signaling Technology (Danvers, MA). All other chemicals were of the analytical grade and obtained from the local chemical suppliers. These chemical reagents were prepared as stock solutions with sterile water (tissue culture grade), and then diluted to the final concentrations before application.

### Cell culture and treatment

Immortalized human lung bronchial epithelial BEAS-2B cells were obtained from the American Type Culture Collection (ATCC, Manassas, VA) and maintained in DMEM supplemented with 10% FBS, 100 U/ml penicillin, and 100 μg/ml streptomycin. The primary human bronchial epithelial cells (NHBEs) were purchased from Lonza (Switzerland) and maintained in BEGM medium (Lonza, Switzerland) supplemented with 100 U/ml penicillin and 100 μg/ml streptomycin. Cells were incubated at 37°C in a humidified atmosphere containing 5% carbon dioxide. In all experiments, cells (0.44 × 10^6^) were seeded into 10 cm-diameter tissue-culture dishes with growth medium and allowed to reach 60% confluence before being exposed to nicotine with doses 0, 500 and 750 μM for 24 hrs, or 0, 10, 25 and 50 μM for 1 to 4 weeks. For competition inhibition assays, 10 μM Roscovitine, 0.1 μM DMAT, 10 or 25 μM LY294002, 10 μM Mecamylamine (Sigma-Aldrich, St. Louis, MO), 10 μM DHβE or 1 μM α-bungarotoxin (MilliporeSigma, St. Louis, MO) was added 1 h before nicotine treatment.

### Cell proliferation assay

The 3-(4,5-dimethylthiazol-2-ul)-2,5-diphenyl tetrasodium bromide (MTT, Sigma-Aldrich, St. Louis, MO) cytotoxicity assay measures mitochondrial reductases, which convert the water soluble MTT salt to a formazan that accumulates in healthy cells. 5000/well cells were seeded into 96-well plate, then cells were exposed to increasing concentrations of nicotine after the cells reached 60% confluency. After 24-hrs of treatment, 10 µL MTT was added to each well containing 100 µL fresh medium with final concentration of 5 mg/mL MTT and incubated for 4 hrs at 37°C in 5% CO_2_. At the end of the incubation, 100 µL of isopropanol containing 0.04 N HCl was added to each well to dissolve the crystals. The absorbance of wells was read at 570 nm using a microplate reader.

### Soft-Agar assays

Cells treated with or without nicotine were rinsed with PBS then seeded in low-gelling temperature Agarose Type VII (Sigma Aldrich, St. Louis, MO). The cells were seeded in triplicate in 6-well plates (5,000 cells/well) in a top layer of 0.35% agarose onto a bottom layer of 0.5% agarose and cultured at 37°C for 6 weeks. Colonies were stained with INT/BCIP (Roche Diagnostics, Indianapolis, IN) overnight for visualization and quantification of colony growth in agar and photographed. Colonies larger than 50 μm were counted. All experiments were performed in triplicates. The results were presented as fold change versus 0 μM nicotine treatment group after the colonies were adjusted for plating efficiency.

### Western blot

Whole cell lysates were prepared from cultured cell using RIPA lysis buffer. For mouse lung tissue lysate, 0.1 g of lung tissue for each sample was shredded and cleaned with pre-cold PBS, followed by sonication in RIPA lysis buffer to get homogenate lysate. Protein concentrations were determined by BCA protein assay (Pierce, Thermo Fisher Scientific, USA). Thirty microgram of whole cell lysate or 80 μg of tissue lysate per lane were separated by 14% SDS-PAGE and blots were transferred to a polyvinylidene difluoride (PVDF) membrane. The membrane was blocked with 5% skimmed milk in TBST for 30 min at room temperature then incubated with primary antibodies overnight at 4°C and secondary antibody conjugated with horseradish peroxidase (HRP) for 1 h at room temperature before visualization by ECL Plus Western blotting detection reagent (Bio-Rad, Hercules, CA). Semi-quantification of the bands were performed using NIH Image J software.

### RNA isolation and Real-time quantitative PCR (RT-qPCR)

Total RNA was extracted using Trizol Reagent (Invitrogen, Grand Island, NY). Briefly, 1 mL Trizol was directly added to the plate to fully digest the cells followed by phase separation and RNA precipitation. For lung tissues, 0.1 g of tissue for each sample was quickly frozen and ground in liquid nitrogen, then mixed with 1 mL Trizol to extract RNA. The quantity and purity of the RNA prepared from each sample were determined using a NanoDrop 2000 spectrophotometer. Reverse transcription (RT) was performed using SuperScript IV First-Strand Synthesis System (Invitrogen, Grand Island, NY) with 1 μg of RNA in a final reaction volume of 20 μL after DNA was removed. After incubation at 50 °C for 50 min, RT was terminated by heating at 85°C for 5 min. Quantitative real-time PCR analysis was performed using Power SYBR Green PCR Master Mix (Applied Biosystems, Waltham, MA). PCR was performed in triplicate. RNA abundance was expressed as 2^−ΔΔCT^ for the target gene normalized against the *GAPDH* gene and presented as fold change to the level in control cells. The following primers were used:

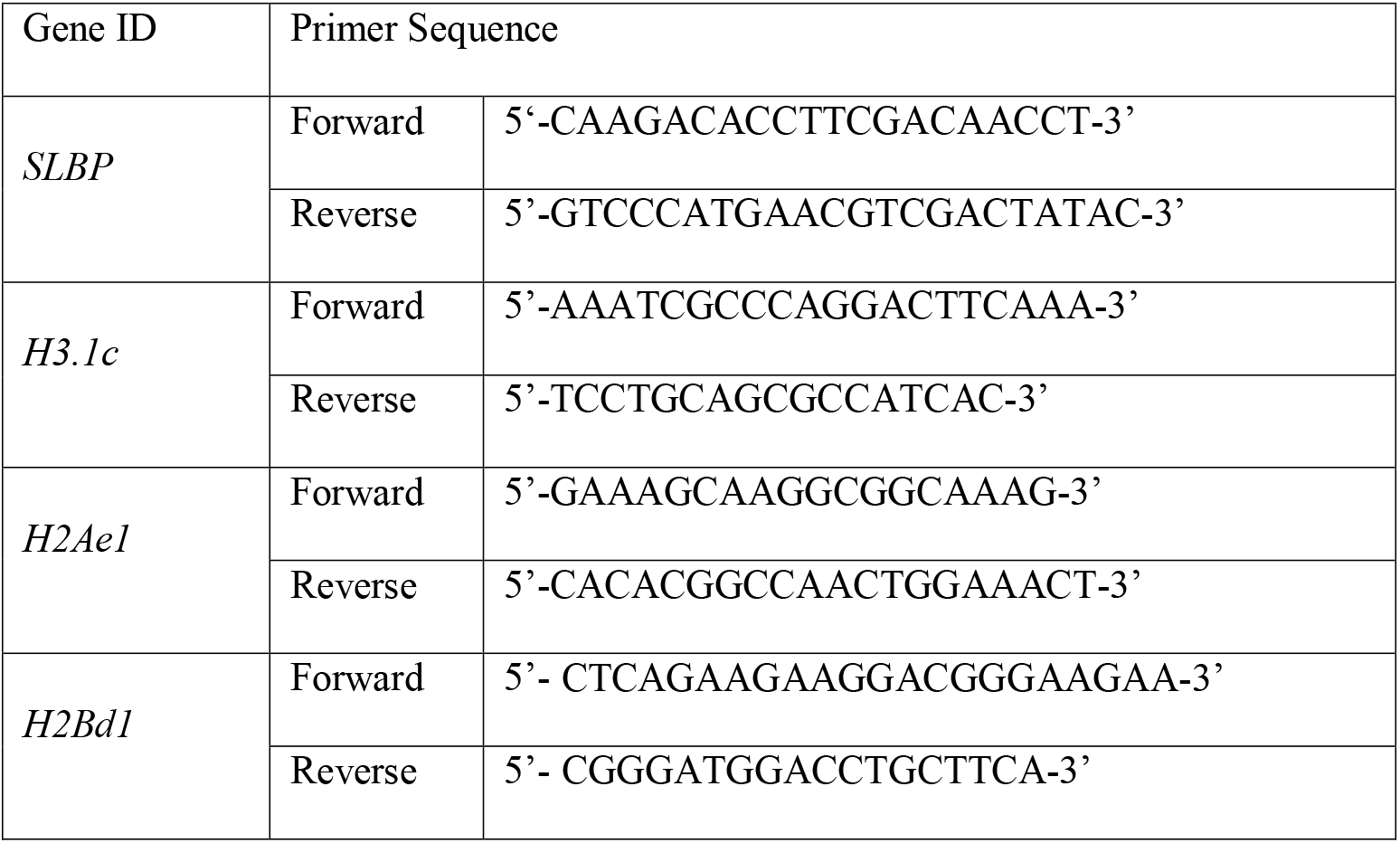
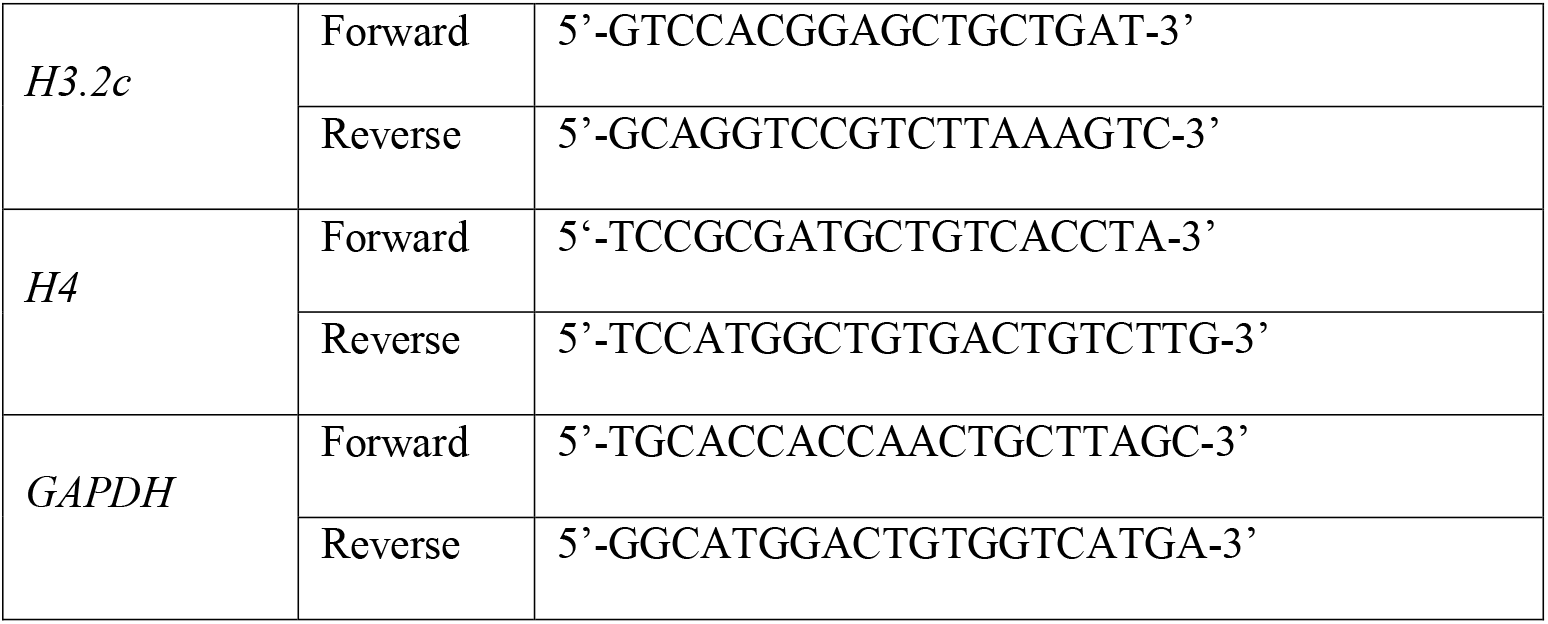

### Immunofluorescence staining

After washing twice with PBS, cells were fixed with 10% formaldehyde, and incubated with 5% BSA at room temperature to block nonspecific binding of antiserum. Cells were incubated with primary antibodies against α7-nAChR (1:150 dilution) at 4 °C overnight. Labeled cells were visualized by the use of FITC-conjugated secondary antibodies (goat anti-rabbit) for 1 hr at room temperature. A final concentration of 0.1 μg/mL DAPI was then added for 10 mins and digital images were captured using ZOE Fluorescent Cell Imager (Bio-Rad). For negative controls, the primary antibodies were omitted.

### Cell transfection and siRNA knockdown

*CDK1, CDK2, CK2*α*1, CK2*α*2*, and α*7-nAChR* siRNAs were purchased from Thermo Fisher (catalog no. 103821, 118638 146139, 103306 and 114074). Cells were seeded 24 hrs before transfection in complete medium without antibiotics to allow 60%-80% confluency at transfection. Lipofectamine RNAiMax (Invitrogen, catalog no. 13778150) and siRNAs were each diluted in serum-free medium. The resulting two solutions were then mixed and incubated to enable complex formation. Cells were then transfected by adding the RNAi-Lipofectamine complex dropwise to medium to achieve a final siRNA concentration of 80 pmol/L (for *CDK1, CDK2, CK2*α*1* and *CK2*α*2*) and 40 pmol/L (for α*7-nAChR*), and cells in negative control group were transfected with universal negative siRNA. Cells were incubated at 37°C and knockdown efficiency was examined at 48 hrs post-transfection.

pcDNA-FLAG-SLBP plasmids were purified using a commercial Qiagen QIAprep Spin Midiprep kit before transfection. Overexpression transfections were performed using Lipofectamine® LTX Reagent with PLUS reagent (Invitrogen, Grand Island, NY) following the manufacturer’s protocol. Briefly, 1.5 × 10^6^ cells were seeded into 6-well dishes 24 hrs prior to transfection. The following day, purified plasmid (1 µg) was transfected into cells in each well with 10 µL of Lipofectamine LTX and 2.5 µL of PLUS reagent. 24 hrs post-transfection, the media was removed and replaced with fresh DMEM. 0.5 µg/mL of G418 selection agent was added to the transfected cells 3 days later. The cells were grown under selection for 3 weeks and harvested for western blot and qPCR analysis.

### E-cig aerosol exposure

Prior to e-cig aerosols exposure, BEAS-2B cells were detached by 0.05% trypsin solution and seeded into Transwell plates containing 0.4 µm pore size polyester membrane inserts in each of 6-well culture plates, at a seeding density of 120,000/insert. NHBE cells were detached by trypsin pre-warmed at 37°C and seeded into the Transwell inserts with at a density of 240,000/insert. Twenty-four hours after seeding, medium on the apical side of the membrane was removed from the insert to create an air-liquid interface (ALI), and cells on the membrane were exposed to either filtered clean air (FA) or e-cig aerosols. Unflavored tobacco e-liquid purchased from a US brand (Halo) was used in this study with 50% PG and 50% VG (50:50 PG/VG), containing 0 or 18 mg/mL nicotine. E-cig aerosols were produced from an automated three-port e-cigarette aerosol generator (e∼Aerosols LLC), as previously described (18). Puff aerosols were generated with charcoal and HEPA-filtered air using a rotor-less diaphragm pump; the puff regimen consisted of 55 mL puff volumes of 3 seconds duration at 30 seconds intervals at 3.6V voltage. Each puff was mixed with filtered dilution air before entering the exposure chamber. After exposure for 20 mins, medium on the apical side of the membrane was added and the cells incubated for 24 hrs in normal culture condition (37 °C in a humidified atmosphere containing 5% carbon dioxide).

### Animal exposure

For *in vivo* experiments, 8-week-old female A/J mice were randomly divided into 3 exposure groups consisting of filtered air (FA) group, PG/VG (50:50) group, and a nicotine (36 mg/ml) plus PG/VG group. The exposure was performed at the University of Rochester. The puff regimen consisted of 55 mL puff volumes of 4 seconds duration at 25 seconds intervals. Mice were exposed to each treatment for 4 hrs per day, 5 days a week, for 3 months. The target exposure was 50 µg/L by filter weight, the flow rate was 100 L/min. Mice were maintained at a temperature of 21-23°C and 40 - 60% humidity. After the final day of exposure, the animals were transferred to the NYU Langone vivarium and analyzed 3 months later, to determine long-term consequences of e-cig exposure. The mice were sacrificed and lung tissues were immediately collected and frozen in −80°C.

### Statistical Analysis

All data were expressed as mean ± standard deviations (SD), significant differences among and between group means were carried out using Software SPSS v22.0 (SPSS Inc., USA). Significant differences between the means were evaluated by analysis of variance test (one-way ANOVA). *Post hoc* tests, when appropriate, were analyzed using an LSD test. Statistical significance was defined as *p* < 0.05.

## Results

### Nicotine exposure induces downregulation of SLBP

Previously we reported that arsenic exposure induces polyadenylation of canonical histone mRNAs through depletion of SLBP, a key factor for pre-mRNA processing of canonical histones (16). Polyadenylation of canonical histone H3.1 mRNA caused aberrant transcription, cell cycle arrest, and chromosome instability in human lung epithelial cells (17). To test the hypothesis that nicotine would have a similar impact on SLBP expression, human lung epithelial BEAS-2B cells were treated with nicotine and the SLBP protein level was measured by western blot analysis.

To determine suitable nicotine concentrations to be used, cell viability MTT assays were performed with BEAS-2B cells treated with either 250 µM, 500 µM, 750 µM or 1000 µM nicotine for 24 h. The doses were chosen because an acute increase of nicotine up to 100 µM in serum and 1000 µM at the mucosal surface immediately following smoking have been reported (19–21). Cell viability was 80% or above at concentrations up to 750 µM (Fig. 1A). As sub-cytotoxic doses (>80% cell viability) are considered appropriate for generating meaningful outcomes in *in vitro* carcinogenesis studies (22,23), cells were treated with either 500 µM or 750 µM. Subsequent experiments at lower doses (∼50 µM) are being carried out to validate some of the observed effects. The SLBP protein levels were reduced by about 25% and 45% following exposures to 500 µM and 750 µM nicotine, respectively, as compared to the control (Fig. 1B). By contrast, *SLBP* mRNA level was decreased only by treatment with 750 µM nicotine, but not by 500 µM, indicating that nicotine downregulates SLBP protein levels perhaps through post-transcriptional regulation at 500 µM (Fig. 1C).

**Figure 1.**
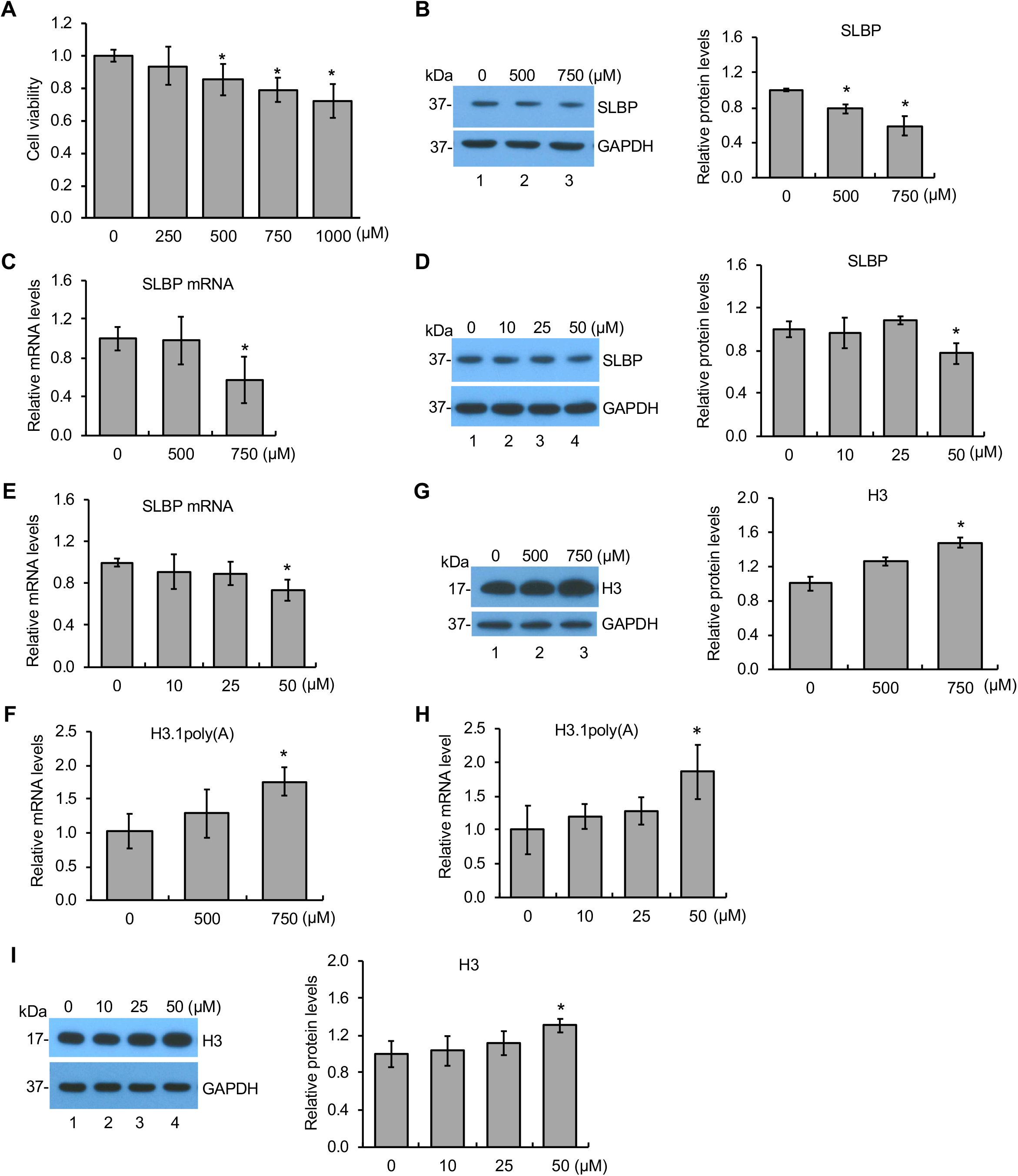
Downregulation of SLBP and polyadenylation of H3.1 mRNA by nicotine. **(A)** BEAS-2B cells were treated with various concentrations of nicotine for 24 hrs and subjected to MTT assays. **(B and C)** Downregulation of SLBP by nicotine. The SLBP protein level and mRNA level were measured by Western blot (B) and RT-qPCR (C), respectively, in BEAS-2B cells treated with or without nicotine for 24 hrs. GAPDH was used as an internal control. The band intensities (left panel in B) were quantified using ImageJ software and presented as bar graphs to show relative quantification of SLBP protein level (right panel in B). **(D and E)** Downregulation of SLBP by low-dose ‘long-term’ treatment with nicotine. The SLBP protein and mRNA levels were measured by Western blot (D) and RT-qPCR (E), respectively, in BEAS-2B cells treated with (10 μM, 25 μM, and 50 μM) or without nicotine for 4 weeks. GAPDH was used as an internal control. The band intensities (left panel in D) were quantified and presented as bar graphs (right panel in D). **(F and G)** Polyadenylation of canonical histone H3.1 mRNA by nicotine. The level of polyadenylated H3.1 mRNA (F) and total H3 protein level (G) were determined by RT-qPCR and Western blot, respectively, in BEAS-2B cells treated with or without nicotine for 24 hrs. The amount of polyadenylated H3.1 mRNA was measured by RT-qPCR using cDNAs synthesized with oligo (dT) primers, capturing polyadenylated mRNAs. GAPDH was used as an internal control. The band intensities (left panel in G) were quantified and presented as bar graphs (right panel in G). **(H and I)** Polyadenylation of canonical histone H3.1 mRNA by low-dose ‘long-term’ treatment with nicotine. The level of polyadenylated H3.1 mRNA (H) and total H3 protein level (I) were determined by RT-qPCR and Western blot, respectively, in BEAS-2B cells treated with (10 μM, 25 μM, and 50 μM) or without nicotine for 4 weeks. GAPDH was used as an internal control. The band intensities (left panel in I) were quantified and presented as bar graphs (right panel in I). Untreated controls were used as references, which were set to 1. The data shown are the mean ± S.D. (n = 3). \**p* < 0.05 vs. untreated control group.

Next, we examined if similar results were observed at lower doses of nicotine that are more relevant to the human scenario. In this case, BEAS-2B cells were treated with 0, 10, 25 and 50 μM of nicotine for 1 to 4 weeks. The SLBP protein levels were decreased starting at 3-weeks during exposure in the 50 μM group (*p* < 0.05) (Fig. 1D and Suppl. Fig. 1A, C and E). The *SLBP* mRNA levels were also decreased following a 4-week treatment with 50 μM of nicotine, as compared to the 0 μM control group (Fig. 1E and Suppl. Fig. 1B, D and F). These data suggest that exposure of the human bronchial epithelial cells to liquid nicotine reduces both protein and mRNA levels for SLBP.

### Nicotine exposure induces polyadenylation of canonical histone H3.1 mRNA and increases H3 protein level

As nicotine exposure reduced SLBP protein level, we investigated whether it induces aberrant 3’-end processing of canonical histone H3.1 mRNA. The level of polyadenylated mRNAs can be determined by using RT-qPCR with oligo (dT) primers for reverse transcription, which captures mRNAs with poly(A) tail. Figure 1F reveals that exposure of BEAS-2B cells to 750 μM of nicotine for 24 hrs results in approximately a 1.5-fold increase of polyadenylated H3.1 mRNA, as compared to the control (Fig. 1F). Polyadenylation of mRNAs for other canonical histones including, H2A, H2B, H3.2, and H4, were also increased by exposure of cells to 750 μM nicotine for 24 hrs (Suppl. Fig. 2A). Given that nicotine exposure induced polyadenylation of all canonical histone mRNAs, we then examined how these alterations changes protein levels of all four core histones. Results of Western blotting showed that the protein levels of total histone H3, as well as H2A, H2B, and H4 were all upregulated following a 24 hr-treatment with 750 μM nicotine, compared to the untreated control group (Fig 1G and Suppl. Fig 2B-E).

Next, we determined if polyadenylation of H3.1 mRNA is also induced by low-dose ‘long-term’ treatment of the cells to nicotine. The level of H3.1 mRNA with a poly(A) tail was significantly increased in BEAS-2B cells treated with 50 μM nicotine for 4 weeks, compared to the 0 μM treatment group (Fig. 1H). In contrast, cells treated either with 50 μM nicotine for 1-3 weeks or with 10 μM or 25 μM nicotine for 4 weeks failed to reach statistical significance, compared to the control group (Fig. 1H and Suppl. Fig. 3A,C and E). The H3 protein level was also upregulated by treatment of cells with 50 μM nicotine for 4-weeks (Fig. 1I and Suppl. Fig. 3B, D and F). These results indicate that *in vitro* exposure to nicotine leads to a loss of SLBP, gain of H3.1 mRNA with poly(A) tail, and an increase in H3 protein.

### Nicotine-mediated SLBP depletion is α7-nAChR dependent

Nicotine is an agonist of nicotinic acetylcholine receptors (nAChRs) (24). To gain mechanistic insight into nicotine-induced SLBP depletion, we asked whether the nicotine-induced downregulation of SLBP requires nAChR activation in BEAS-2B cells. nAChRs belong to the superfamily of homologous Cys-loop ion channel receptors, consisting of nine α (α2 to α10) and three β subunits (β2-β4). The upregulation of α7-nAChR has been shown to be particularly important in lung cancer, facilitating the proliferation and migration of cancer cells in lung tissue (25,26). Immunofluorescence staining of BEAS-2B cells demonstrated a dose-dependent increase of α7-nAChR protein upon nicotine treatment (Fig. 2A). α-BTX, a classical α7-nAChR inhibitor, completely abrogated nicotine-induced upregulation of α7-nAChR (Fig. 2A). Western blot analysis confirmed the increase of α7-nAChR protein levels by nicotine treatment (Fig. 2B and C). The nicotine-induced upregulation of α7-nAChR protein was suppressed by α-BTX, but not by DHβE, an inhibitor of α3/α4 subunits of nAChRs (Fig. 2B and C). Importantly, nicotine-induced reduction of SLBP protein level was attenuated by α7-nAChR inhibitor α-BTX but not by DHβE (Fig. 2B and D). This finding suggests that the downregulation of SLBP by nicotine is likely dependent specifically on α7-nAChR.

**Figure 2.**
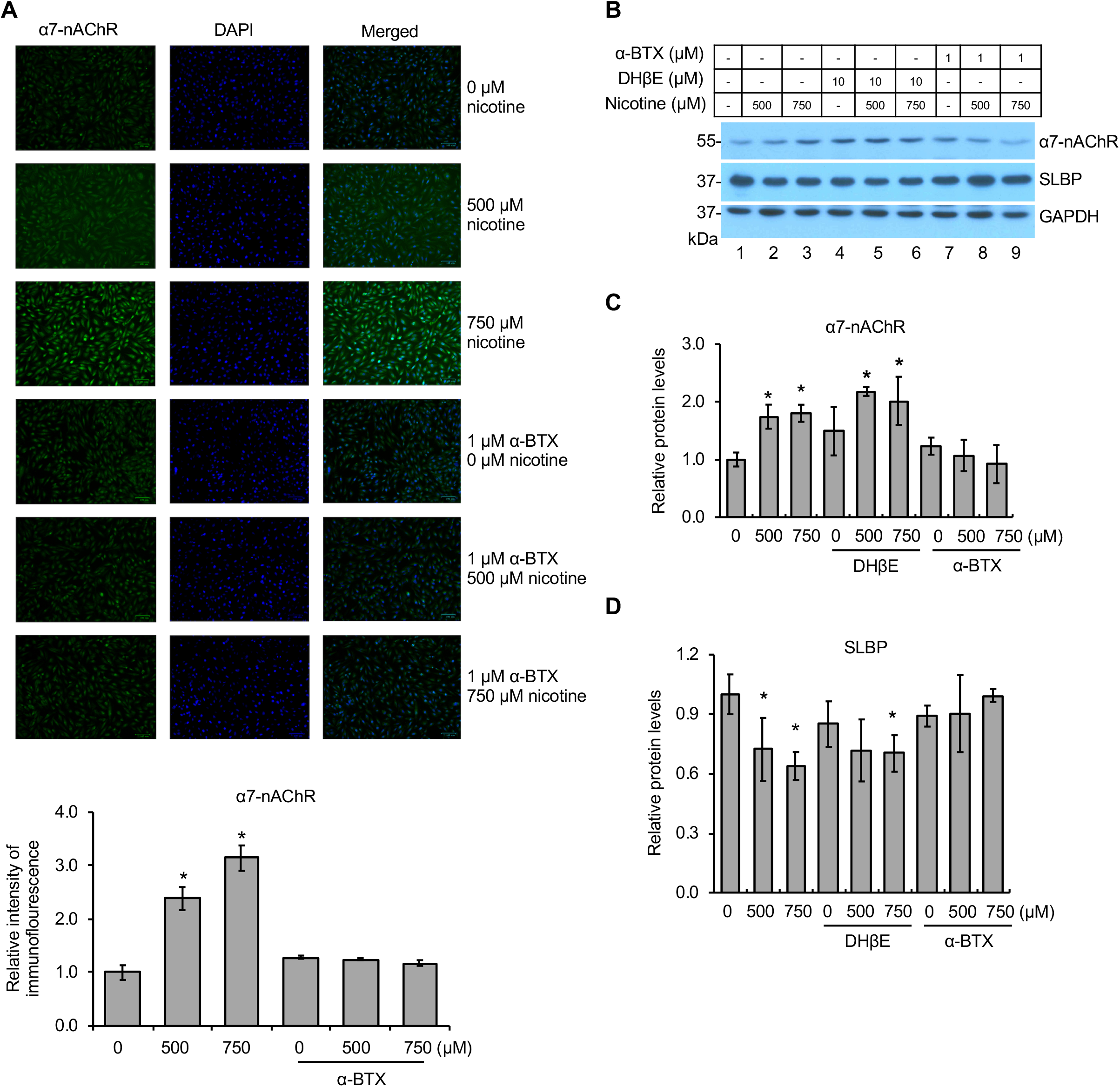
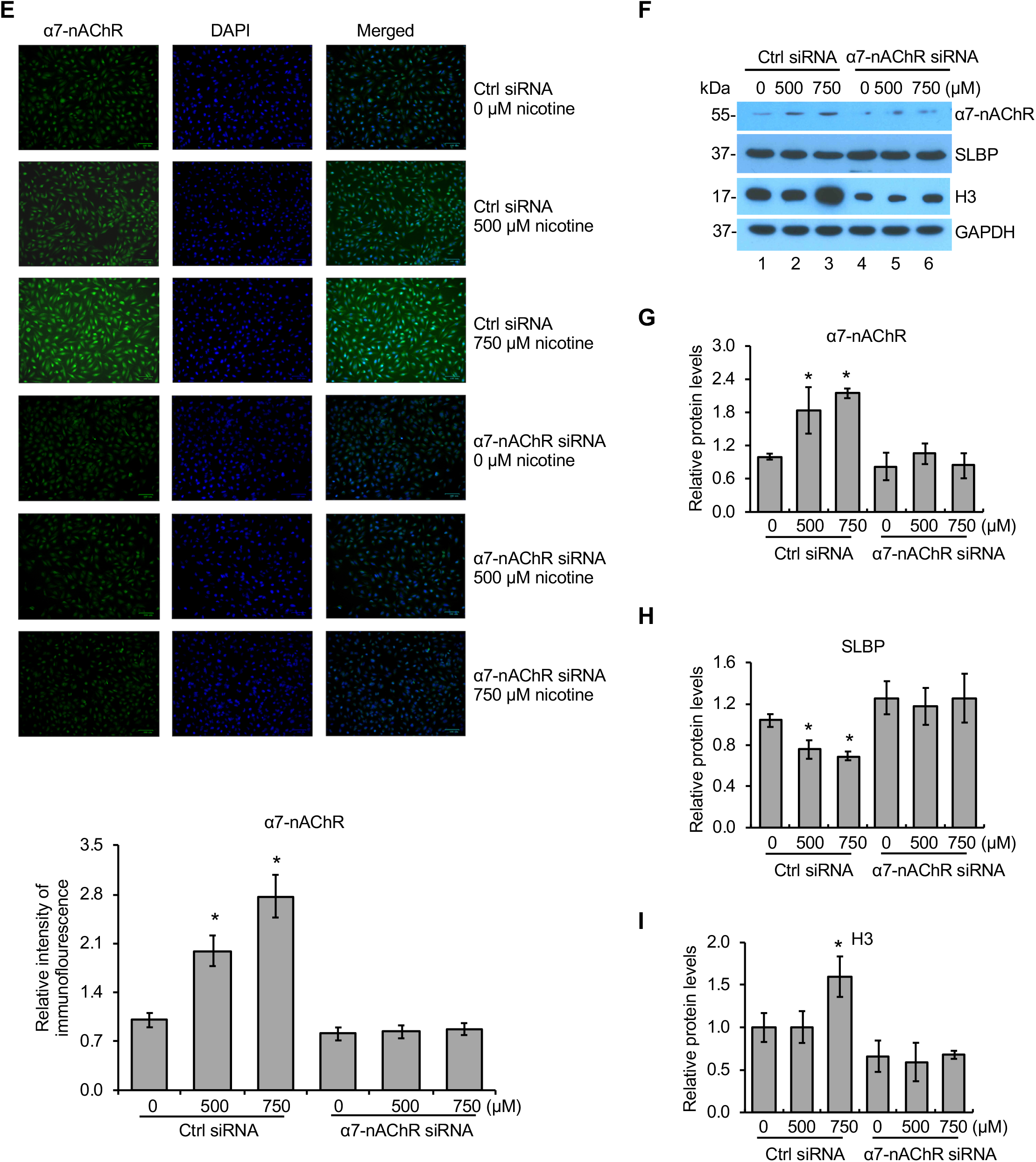
Nicotine-mediated downregulation of SLBP is a7-nAChR dependent. **(A)** Immunofluorescence (IF) staining of α7-nAChR in BEAS-2B cells treated with or without nicotine for 24 hrs in the presence or absence of 1 μM α-BTX, an inhibitor of α7-nAChR. Representative IF staining for α7-nAChR (green), DAPI (blue), and merged images are shown. The IF intensities for α7-nAChR were determined by ImageJ software, normalized to control group, and presented as bar graphs (bottom panel). **(B-D)** α-BTX, an inhibitor of α7-nAChR, but not DHβE, an inhibitor of α3/4-nAChR, attenuates nicotine-induced downregulation of SLBP. BEAS-2B cells were treated with or without nicotine for 24 hrs in the presence or absence of DHβE or α-BTX and subjected to western blot analysis with indicated antibodies (B). The band intensities were quantified and presented as bar graphs to show relative protein levels for α7-nAChR (C) and SLBP (D). GAPDH was used as an internal control. **(E)** IF staining of α7-nAChR following nicotine treatment of BEAS-2B cells transiently transfected with the control siRNA or the siRNA specific for α7-nAChR. Representative IF staining for α7-nAChR (green), DAPI (blue), and merged images are shown. The IF intensities for α7-nAChR were determined by ImageJ software, normalized to control group, and presented as bar graphs (bottom panel). **(F-I)** Knockdown of α7-nAChR by siRNA attenuates nicotine-mediated downregulation of SLBP. BEAS-2B cells that have been transiently transfected with the control siRNA or the siRNA specific for α7-nAChR were treated with or without nicotine for 24 hrs and subjected to western blot analysis with indicated antibodies (F). The band intensities were quantified and presented as bar graphs to show relative protein levels for α7-nAChR (G), SLBP (H), and total H3 (I). GAPDH was used as an internal control. Untreated controls in lane 1 were used as references, which were set to 1. The data shown are the mean ± S.D. (n = 3). \**p* < 0.05 vs. untreated control group.

To further validate an essential role for α7-nAChR in nicotine-induced SLBP depletion, siRNA was used to knock down α7-nAChR expression. Both immunofluorescence staining and western blot analysis revealed a dose-dependent increase of α7-nAChR protein expression in nicotine-treated BEAS-2B cells that were transfected with the control siRNAs. In contrast, transfection of siRNA specific for α7-nAChR abrogated the nicotine-mediated upregulation of α7-nAChR (Fig. 2E-G). Notably, nicotine-induced depletion of SLBP and upregulation of H3 protein levels were attenuated by knockdown of α7-nAChR expression by siRNA (Fig. 2F, H and I). Together, we conclude that the nicotine-induced decrease in SLBP and consequent increase in H3 are α7-nAChR-dependent.

### Downregulation of SLBP *via* PI3K/AKT signal transduction pathway

It has been reported that the stimulation of nAChRs activates the PI3K/AKT pathway (27). As nicotine upregulated α7-nAChR, we next investigated if PI3K/AKT pathway is mechanistically involved in the nicotine-induced SLBP depletion. As shown in Figure 3, nicotine treatment significantly increased phosphorylation of AKT at S473 (p-AKT^S473^), but not p-AKT ^T308^, measured either directly or relative to total AKT (Fig. 3A and B). Total AKT protein levels were not changed by nicotine treatment. To determine whether PI3K activation was involved in the depletion of SLBP, cells were pretreated with 10 μM or 25 μM LY294002, an inhibitor of PI3K, prior to exposure of the cells to nicotine. In control cells, the SLBP protein level was significantly downregulated following nicotine treatment (Fig. 3C and D, lanes 1-3). Whereas the SLBP protein level was slightly (albeit, not significantly) decreased in the presence of 10 μM LY294002, the nicotine-induced SLBP reduction was apparently rescued by 25 μM LY294002 (Fig. 3C and D, lanes 4-9). The H3 protein level was unchanged by nicotine treatment in the presence of either 10 μM or 25 μM LY294002 (Fig. 3C and E). These results suggest that the activation of PI3K/AKT pathway is involved in the nicotine-induced downregulation of SLBP.

**Figure 3.**
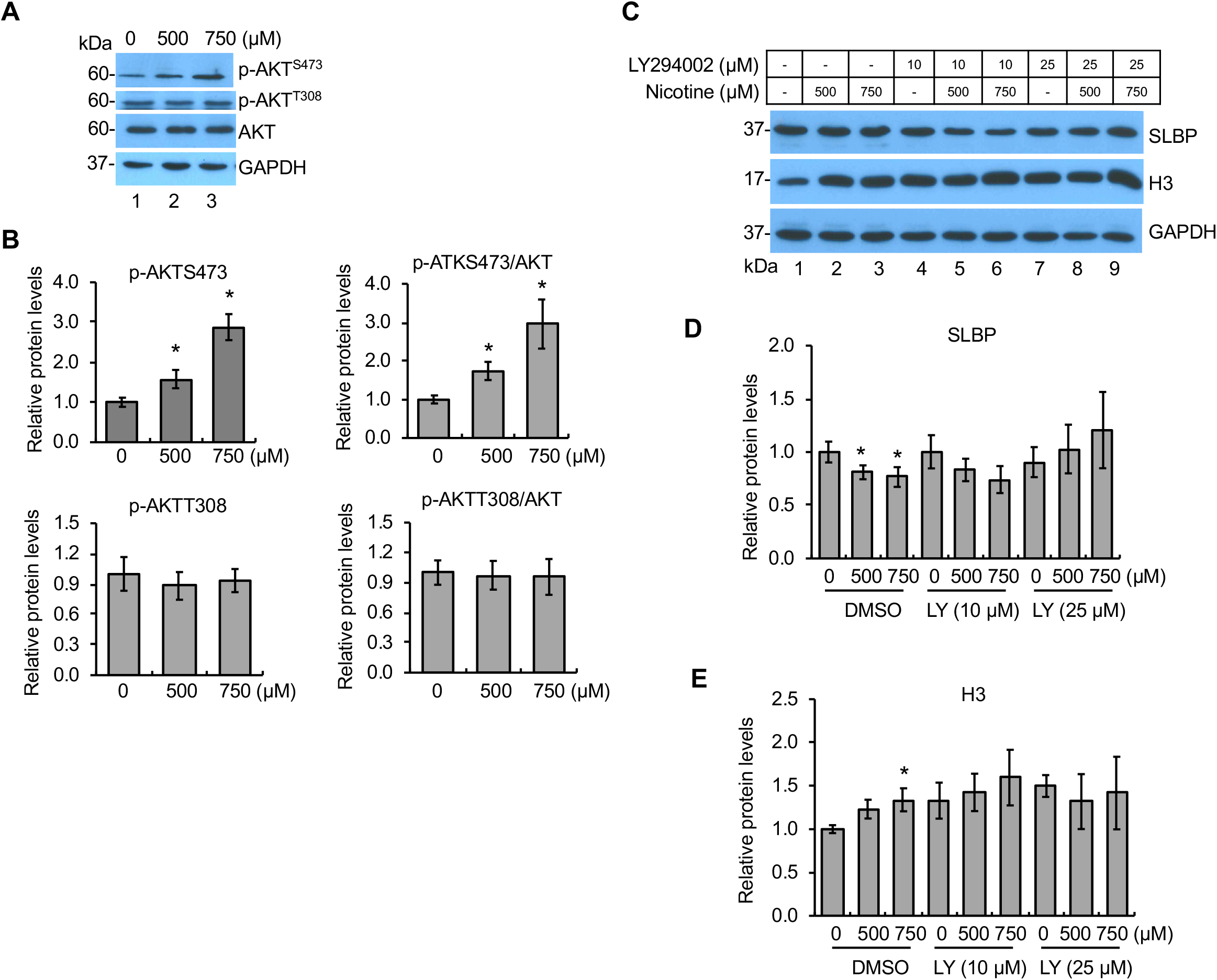
Involvement of PI3K/AKT pathway in nicotine-induced downregulation of SLBP. **(A and B)** Phosphorylation of AKT at S473 by nicotine. BEAS-2B cells were treated with or without nicotine for 24 hrs followed by western blot analysis with indicated antibodies (A). The band intensities were quantified and presented as bar graphs (B). GAPDH was used as an internal control. **(C-E)** Inhibition of PI3K/AKT pathway attenuates nicotine-induced downregulation of SLBP. BEAS-2B cells were treated with or without nicotine for 24 hrs in the presence or absence of LY294002, an inhibitor of PI3K, and subjected to Western blot with indicated antibodies (C). The band intensities were quantified and presented as bar graphs to show relative quantifications of SLBP (D) and total H3 (E). GAPDH was used as an internal control. Untreated controls in lane 1 were used as references, which were set to 1. The data shown are the mean ± S.D. (n = 3). \**p* < 0.05 vs. untreated control group.

### SLBP depletion by nicotine is regulated by CDK1/2

The level of SLBP is mainly regulated by post-transcriptional mechanisms in normal cells (28). Previous studies have demonstrated that phosphorylation of SLBP at Thr61 by cyclin A/CDK1 primes phosphorylation at Thr60 by CK2, which triggers subsequent proteasome-mediated SLBP degradation at the S/G2 cell cycle border (29). Thus, we examined if CDK1 and/or CK2 are involved in nicotine-induced SLBP depletion. We first measured the changes of SLBP protein levels following nicotine treatment in the presence of roscovitine, an inhibitor of CDK1/2. Nicotine exposure reduced the SLBP protein level with statistical significance in the absence of roscovitine (compare lane 1 with lanes 3 and 4 in Fig. 4A and Fig. 4B), a similar reduction was not observed in the cells pretreated with roscovitine (compare lane 2 with lanes 5-6 in Fig. 4A and Fig. 4B), suggesting that CDK1/2 is required for nicotine-induced SLBP depletion. Meanwhile, the upregulation of H3 protein level by nicotine was attenuated in the presence of roscovitine (Fig. 4A and C).

**Figure 4.**
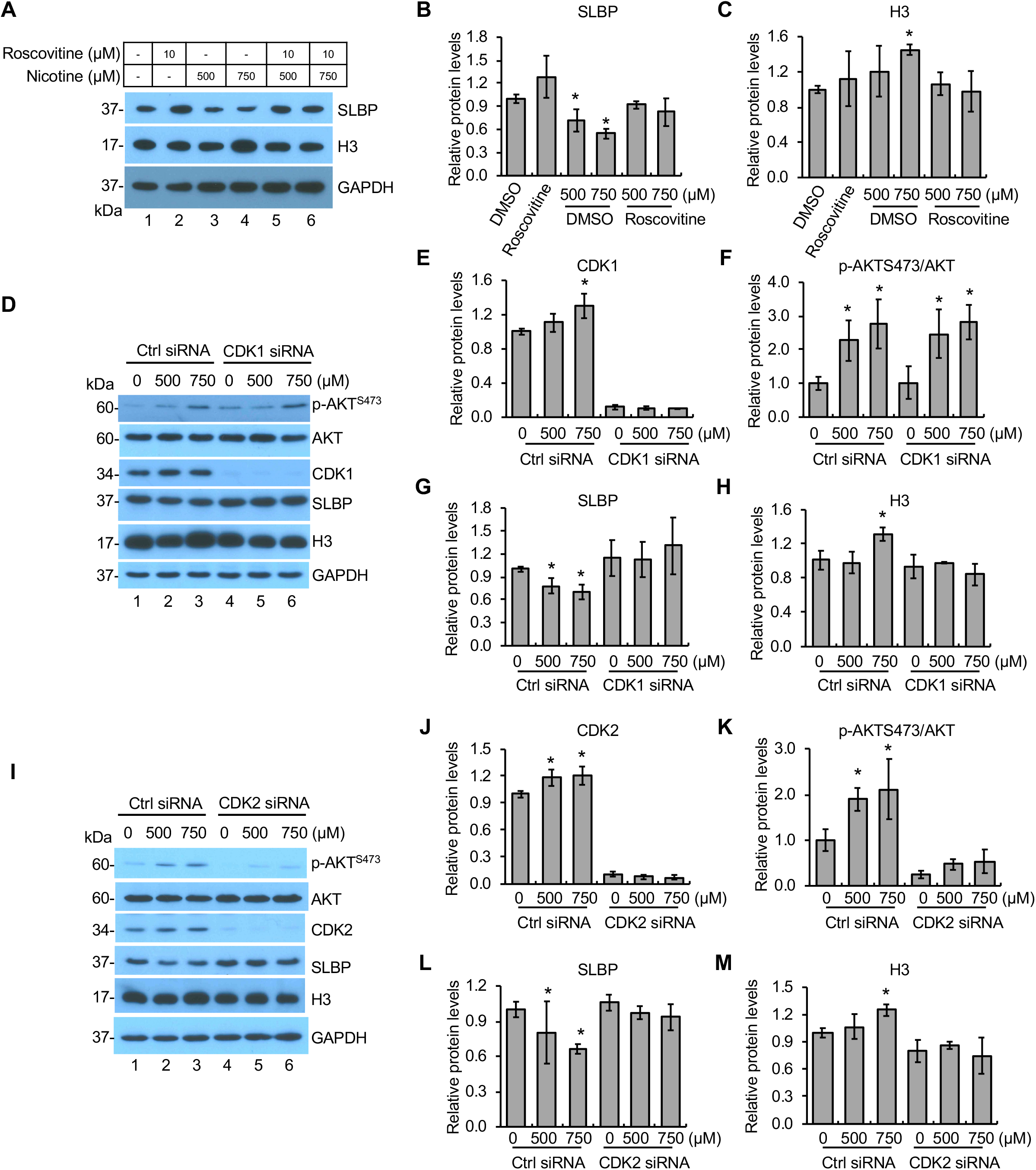
Nicotine-induced downregulation of SLBP is regulated by CDK1/2. **(A-C)** Inhibition of CDKs attenuates nicotine-induced downregulation of SLBP. BEAS-2B cells were treated with or without nicotine for 24 hrs in the presence or absence of roscovitine, an CDK inhibitor, and subjected to Western blot (A). The band intensities were quantified and presented as bar graphs to show relative quantifications of SLBP (B) and H3 (C). GAPDH was used as an internal control. **(D-H)** Knockdown of CDK1 by siRNA attenuates nicotine-induced downregulation of SLBP but not phosphorylation of AKT at S473. BEAS-2B cells were transiently transfected with control siRNA or *CDK1* siRNA and then treated with or without nicotine for 24 hrs followed by western blot analysis with indicated antibodies (D). The band intensities were quantified and presented as bar graphs to show relative quantifications of CDK1 (E), p-AKTS473/AKT (F), SLBP (G), and total H3 (H). **(I-M)** Knockdown of CDK2 by siRNA attenuates nicotine-induced downregulation of SLBP and phosphorylation of AKT at S473. BEAS-2B cells were transiently transfected with control siRNA or *CDK2* siRNA and then treated with or without nicotine for 24 hrs followed by western blot analysis with indicated antibodies (I). The band intensities were quantified and presented as bar graphs to show relative quantifications of CDK2 (J), p-AKTS473/AKT (K), SLBP (L), and total H3 (M). Untreated controls in lane 1 were used as references, which were set to 1. The data shown are the mean ± S.D. (n = 3). \**p* < 0.05 vs. untreated control group.

For further verifying the role of CDK1/2 in inducing nicotine-induced SLBP downregulation, BEAS-2B cells were transfected with control siRNA or siRNAs specific for either *CDK1* or *CDK2* followed by the western blot analysis. The results showed highly efficient knockdown of *CDK1* or *CDK2* expressions by respective siRNAs (Fig. 4D and I and Fig. 4E and J). In the control siRNA cells, the levels of both CDK1 and CDK2 were slightly increased following nicotine treatment (Lanes 1-3 in Fig. 4D and I; Fig. 4E and J). p-AKT^S473^ was upregulated by nicotine exposure in the control cells as seen in the ‘wild-type’ BEAS-2B cells (Lanes 1-3 in Fig. 4D and I; Fig. 4F and K). Interestingly, *CDK1* siRNA had no effect on nicotine-induced changes in p-AKT^S473^(Fig. 4D and F). By contrast, the level of p-AKT^S473^ was greatly inhibited in the *CDK2* siRNA cells (Fig. 4I and K), suggesting that CDK2 but not CDK1 was responsible for upregulation of p-AKT^S473^ by nicotine exposure. However, knockdown of both CDK1 and CDK2 was able to reverse nicotine-induced depletion of SLBP (Fig. 4D and G; Fig. 4I and L) and upregulation of histone H3 (Fig. 4D and H; Fig 4I and M), indicating that both CDK1 and CDK2 are involved in nicotine-mediated SLBP depletion likely through different mechanisms that are either p-AKT^S473^-dependent (CDK2) or independent (CDK1).

### SLBP depletion by nicotine is regulated by CK2

To test if CK2 is also involved in nicotine-mediated SLBP downregulation, we investigated how DMAT, an inhibitor of CK2, affects SLBP protein level following nicotine exposure in BEAS-2B cells. In the absence of DMAT, the SLBP protein level was reduced dose-dependently by nicotine treatment (lanes 1-3 in Fig. 5A and B), whereas the SLBP depletion was rescued in the presence of DMAT (lanes 4-6 in Fig. 5A and B). Additionally, the western blotting results showed that pretreatment of the cells with DMAT prevented nicotine-induced upregulation of H3 protein level (Fig. 5A and C). These data suggest a role for CK2 might in the nicotine-induced loss of SLBP.

**Figure 5.**
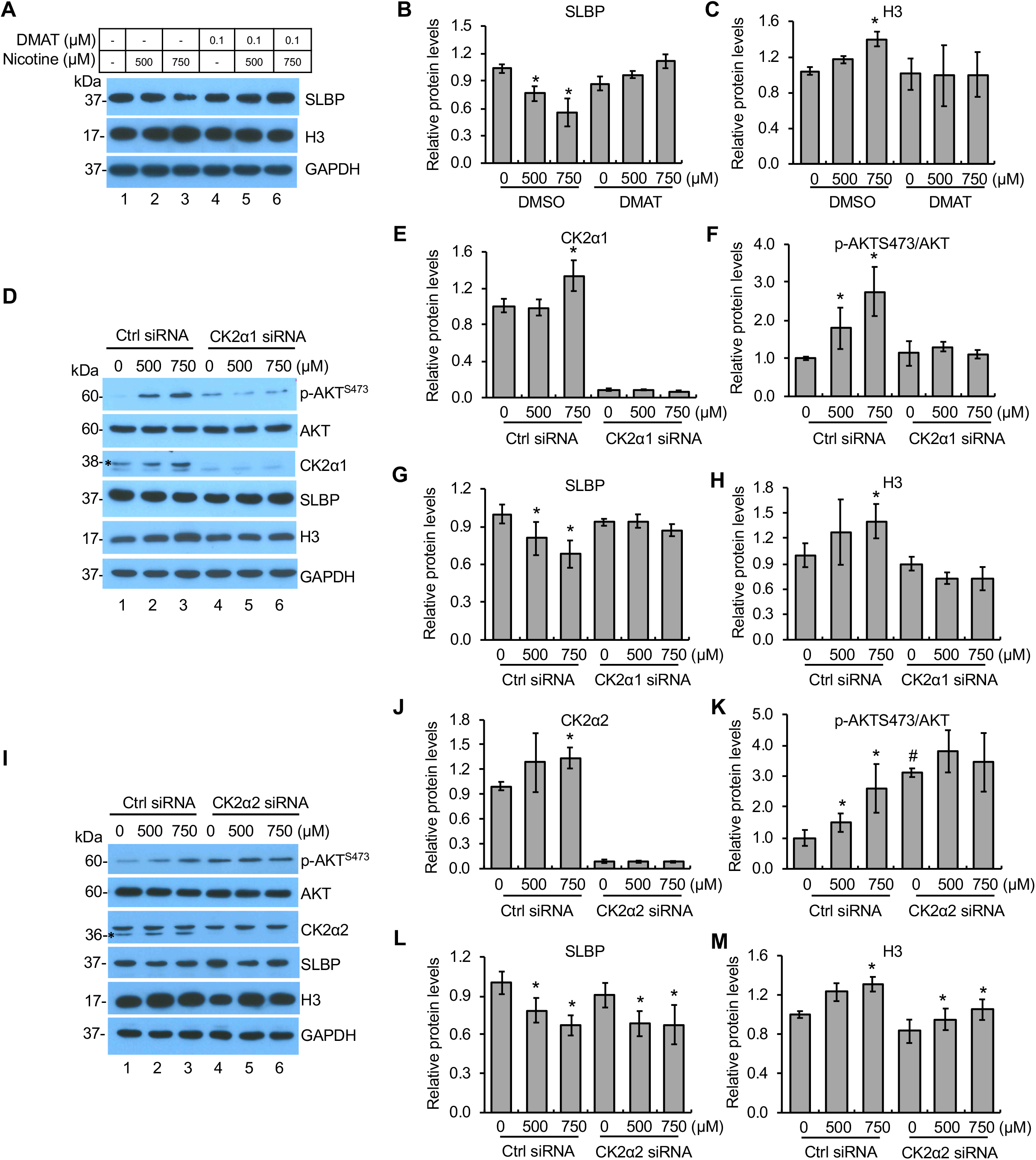
Nicotine-induced downregulation of SLBP is regulated by CK2. **(A-C)** Inhibition of CK2 attenuates nicotine-induced downregulation of SLBP. BEAS-2B cells were treated with or without nicotine for 24 hrs in the presence or absence of DMAT, an CK2 inhibitor, and subjected to Western blot (A). The band intensities were quantified and presented as bar graphs to show relative quantifications of SLBP (B) and H3 (C). GAPDH was used as an internal control. **(D-H)** Knockdown of CK2α1 by siRNA attenuates nicotine-induced downregulation of SLBP and phosphorylation of AKT at S473. BEAS-2B cells were transiently transfected with control siRNA or *CK2*α*1* siRNA and then treated with or without nicotine for 24 hrs followed by western blot analysis with indicated antibodies (D). The band intensities were quantified and presented as bar graphs to show relative quantifications of CK2α1 (E), p-AKTS473/AKT (F), SLBP (G), and total H3 (H). GAPDH was used as an internal control. **(I-M)** Knockdown of CK2α2 by siRNA has no effects on nicotine-induced downregulation of SLBP. BEAS-2B cells were transiently transfected with control siRNA or *CK2*α*2* siRNA and then treated with or without nicotine for 24 hrs followed by western blot analysis with indicated antibodies (I). The band intensities were quantified and presented as bar graphs to show relative quantifications of CK2α2 (J), p-AKTS473/AKT (K), SLBP (L), and total H3 (M). GAPDH was used as an internal control. Untreated controls in lane 1 were used as references, which were set to 1. The data shown are the mean ± S.D. (n = 3). \**p* < 0.05 vs. untreated control group; ^#^*p* < 0.05 vs. control group in Ctrl siRNA cells in panel K.

The role of CK2 in SLBP downregulation was further verified by using siRNAs targeting the α1 and α2 catalytic subunit, respectively. Both siRNAs specifically depleted the expression of respective targeting subunits (compare lane 1 and 4 in Fig. 5D and I; Fig. 5E and J). The protein levels for CK2α1 and CK2α2 were upregulated by nicotine treatment in the control siRNA cells (lanes 1-3 in Fig. 5D and I; Fig. 5E and J), while the induction was not observed in the *CK2*α*1*- and *CK2*α2-siRNA cells (lanes 4-6 in Fig. 5D and I; Fig.5E and J). Notably, the knockdown of CK2α1 reversed nicotine-induced p-AKT^S473^, the loss of SLBP and the gain of histone H3 protein (lanes 4-6 in Fig. 5D; Fig. 5F-H). By contrast, the CK2α2 knockdown failed to reverse the depletion of SLBP and the upregulation of H3 protein levels induced by nicotine exposure (lanes 4-6 in Fig. 5I; Fig. 5L and M). However, the level of p-AKT^S473^ was not further increased by nicotine treatment in the CK2α2 siRNA cells probably due to the amount of p-AKT^S473^ has already reached saturation by CK2α2 knockdown even prior to the nicotine treatment (Fig. 5I and K). Together, these results suggest that the α1 catalytic subunit but not the α2 subunit of CK2 is necessary for nicotine-induced SLBP depletion.

### Nicotine activates CDK1/2, CK2, and AKT *via* α7-nAChR

We demonstrated that nicotine exposure activates CDK1/2, CK2, and AKT, which are required for nicotine-induced downregulation of SLBP. To determine if these changes induced by nicotine are α7-nAChR-dependent, we performed western blot analysis with the cells transfected with the α*7-nAChR* siRNA or the control siRNA. In the control cells, α7-nAChR protein level was increased by about 2-fold following nicotine treatment, whereas no increase was observed in the cells transfected with siRNA for α*7-nAChR*, indicating that induction of α7-nAChR was successfully inhibited by the α7-nAChR-specific siRNA (Fig. 6A and B). Consistent with the results obtained with the ‘wild-type’ original BEAS-2B cells, the levels of p-AKT^S473^, CDK1/2, and CK2 were significantly increased following nicotine treatment in the BEAS-2B cells transfected with control siRNA cells, best revealed with 750 μM nicotine (lanes 1-3 in Fig. 6A; Fig. 6C and F-H). As expected, nicotine treatment resulted in the reduction of SLBP and the increase in H3 protein level in the siRNA control cells (Fig. 6A, D and E). Importantly, the increases in p-AKT^S473^ and CDK1/2 and CK2 protein levels mediated by nicotine exposure were prevented by the knockdown of α7-nAChR (lanes 4-6 in Fig. 6A; Fig. 6C and F-H). These data suggest that nicotine-induced phosphorylation of AKT at S473 and upregulation of CDK1/2 and CK2 are α7-nAChR-dependent.

**Figure 6.**
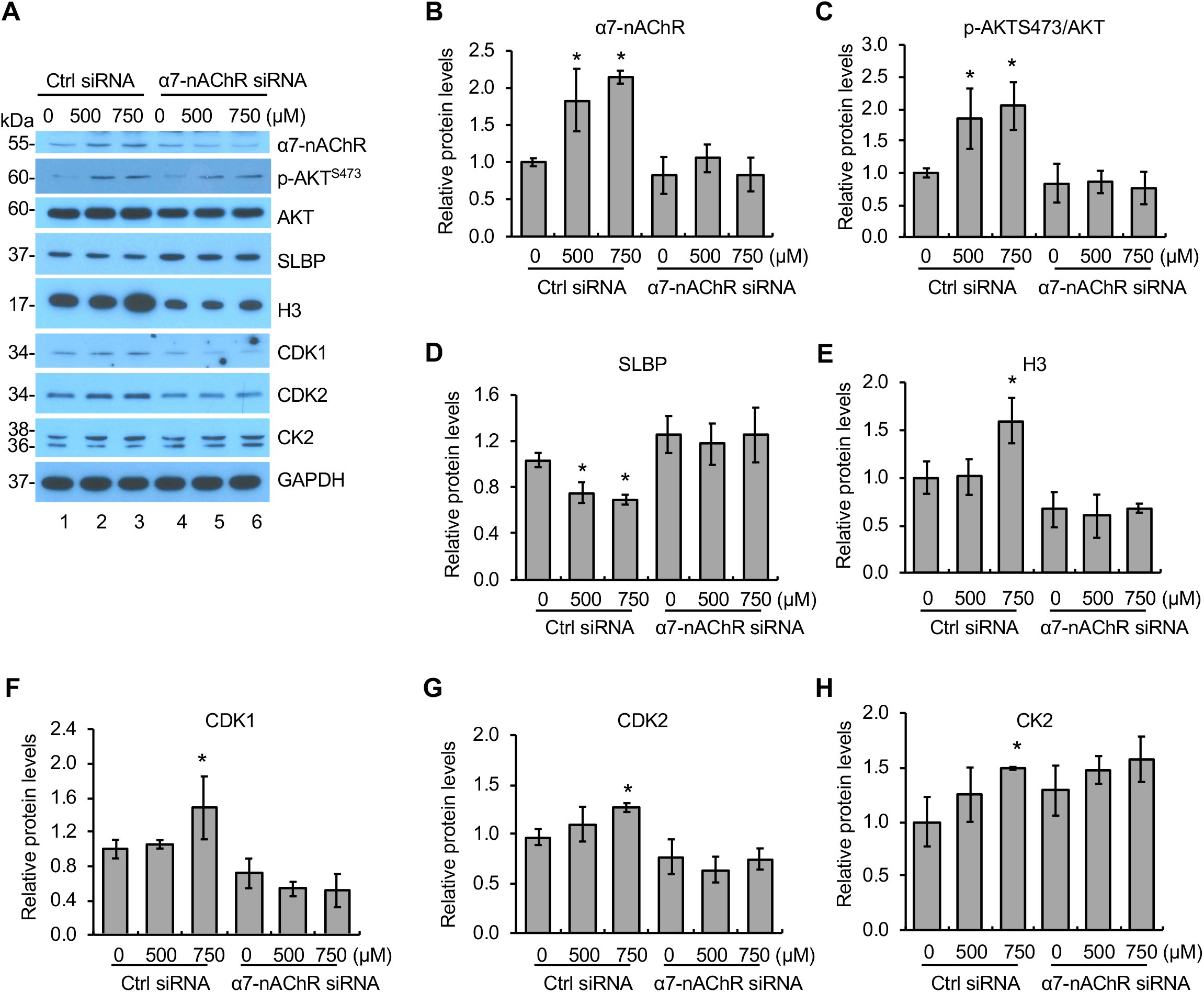
Nicotine-induced activation of AKT, CDK1/2, and CK2 is α7-nAChR dependent. **(A)** Knockdown of α7-nAChR by siRNA attenuates nicotine-induced activation of AKT, CDK1/2, and CK2. BEAS-2B cells were transiently transfected with control siRNA or α*7-nAChR* siRNA and then treated with or without nicotine for 24 hrs followed by western blot analysis with indicated antibodies. **(B-H)** The band intensities from (A) were quantified and presented as bar graphs to show relative quantifications of α7-nAChR (B), p-AKTS473/AKT (C), SLBP (D), total H3 (E), CDK1 (F), CDK2 (G), and CK2 (H). GAPDH was used as an internal control. The controls in lane 1 were used as references. The data shown are the mean ± S.D. (n = 3). \**p* < 0.05 vs. control group.

### Nicotine-induced cell transformation is α7-nAChR-dependent

To explore a potential role for α7-nAChR in nicotine-induced cell transformation, we carried out a series of soft agar assays, which measure the ability of cells to grow anchorage independently. We first examined if nicotine can induce cell transformation. Treatment of BEAS-2B cells with 750 μM of nicotine for 24 hr facilitated colony formation in soft agar as compared to the untreated control cells (Fig. 7A). We also examined if ‘long-term’ low-dose nicotine treatment could induce anchorage-independent growth of BEAS-2B cells. The cells were treated with 10, 25 and 50 μM of nicotine for 1 to 4 weeks. Exposure of BEAS-2B cells with 50 μM nicotine for 4 weeks enhanced colony formation in soft agar with a statistical significance as compared to the 0 μM nicotine treatment group (Fig. 7B and Suppl. Fig. 4). These results indicate that nicotine is able to induce cell transformation *in vitro*.

**Figure 7.**
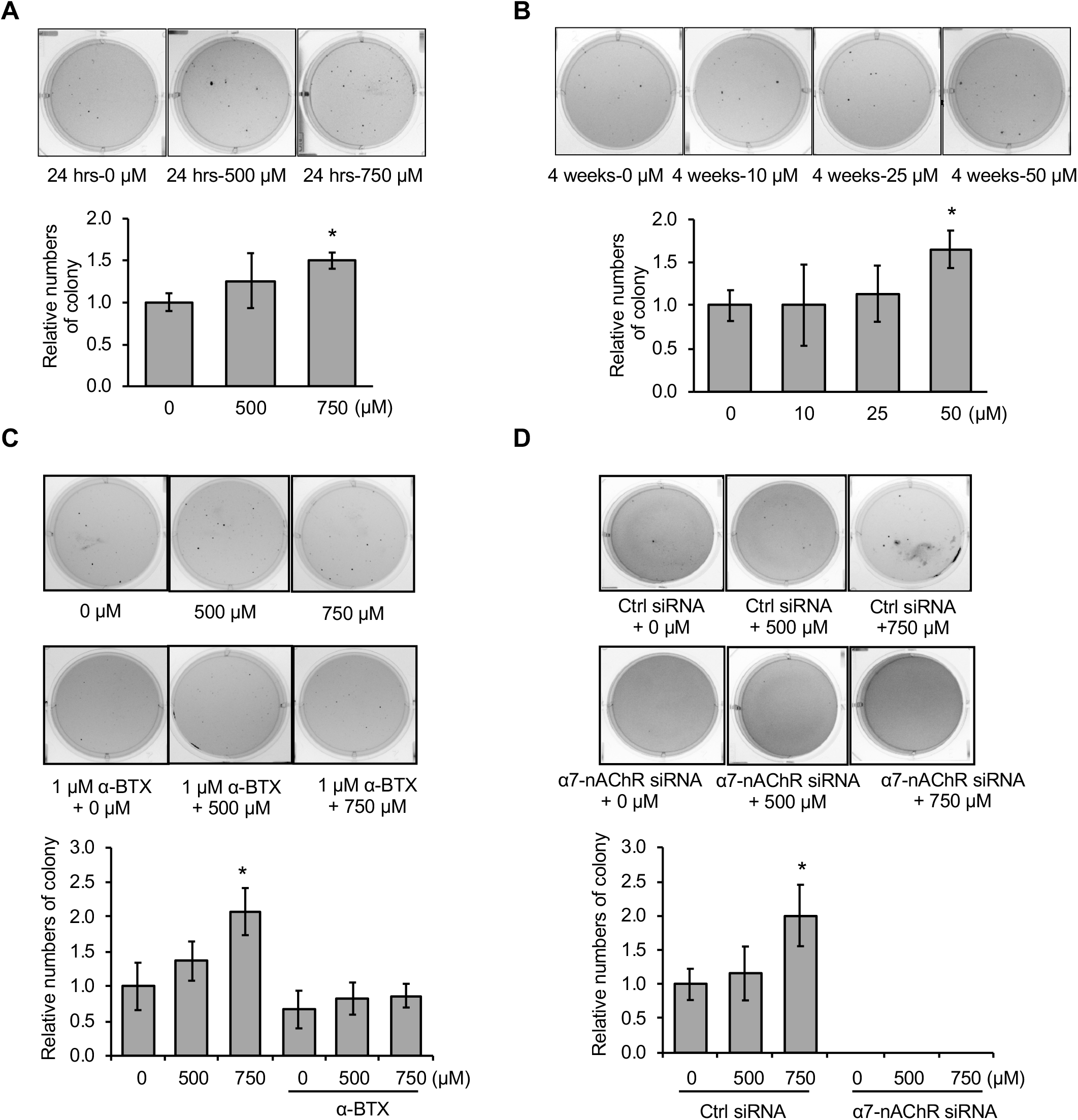
Nicotine-induced cell transformation is α7-nAChR-dependent. **(A and B)** Exposure of BEAS-2B cells to 750 μM nicotine for 24 hrs (A) or 50 μM for 4 weeks (B) enhances anchorage-independent cell growth. After nicotine treatment, the cells were plated in soft agar, and cultured for 6 weeks. The data shown are the mean ± S.D. (n = 3). \**p* < 0.05. vs. control group. **(C)** Inhibition of α7-nAChR attenuates nicotine-induced anchorage-independent cell growth. BEAS-2B cells were treated with or without nicotine for 24 hrs in the presence or absence of α-BTX, an inhibitor of α7-nAChR, and then subjected to soft agar assays. The cells were grown in soft agar for 6 weeks. The data shown are the mean ± S.D. (n = 3). \**p* < 0.05 vs. control group. **(D)** Knockdown of α7-nAChR inhibits anchorage-independent cell growth. BEAS-2B cells were transiently transfected with control siRNA or siRNA specific for α*7-nAChR*, treated with or without nicotine for 24 hrs, and then subjected to soft agar assays. The cells were grown in soft agar for 6 weeks. The data shown are the mean ± S.D. (n = 3). \**p* < 0.05 vs. control group.

We then treated BEAS-2B cells with α-BTX, an inhibitor of α7-nAChR, or transfected the cells with siRNA for α*7-nAChR*, prior to exposure of the cells to nicotine, and then performed soft-agar assays. Nicotine-mediated colony formation in soft agar was suppressed by pretreatment of the cells with α-BTX (Fig. 7C). Moreover, knockdown of α7-nAChR by siRNA abolished colony formation in soft agar, regardless of whether the cells were treated with or without nicotine (Fig. 7D), suggesting that α7-nAChR is required for anchorage-independent cell growth. Taken together, we conclude that nicotine induces cell transformation through the α7-nAChR.

### SLBP overexpression inhibits nicotine-induced cell transformation

Nicotine exposure induced cell transformation *via* α7-nAChR. Since nicotine exposure led to SLBP protein reduction and H3 protein upregulation *via* α7-nAChR, we explored the role of SLBP depletion in nicotine-induced cell transformation. To this end, we established BEAS-2B cell lines that stably express FLAG-tagged SLBP. We chose two SLBP-overexpressing cell clones (SLBP clone 1 and clone 2) that expressed relatively low level of exogenous SLBP (Fig. 8A) to determine how the overexpression of SLBP affects nicotine-induced cell transformation. BEAS-2B cells and SLBP clone 1 and clone 2 were treated with either 0, 500 or 750 μM nicotine for 24 hr. In parental BEAS-2B cells, nicotine downregulated SLBP and upregulated polyadenylated H3.1 mRNA and H3 protein (Fig. 8A-E). In contrast, overexpression of SLBP in both SLBP clone 1 and 2 prevented nicotine-induced loss of SLBP and increase of polyadenylated H3.1 mRNA and H3 protein (Fig. 8A-E). Importantly, while colony numbers in soft agars were increased with statistical significance by treatment of the parental cells with 750 μM nicotine for 24 hrs, cells that expressed FLAG-SLBP exhibited no colony formation in soft agar in the presence or absence of nicotine treatment (Fig. 8F). Similar results were obtained when the cells were treated with 50 μM nicotine for 4 weeks (Fig. 8G). These results suggest that SLBP has an inhibitory effect on anchorage-independent cell growth and is likely required for nicotine-induced cell transformation.

**Figure 8.**
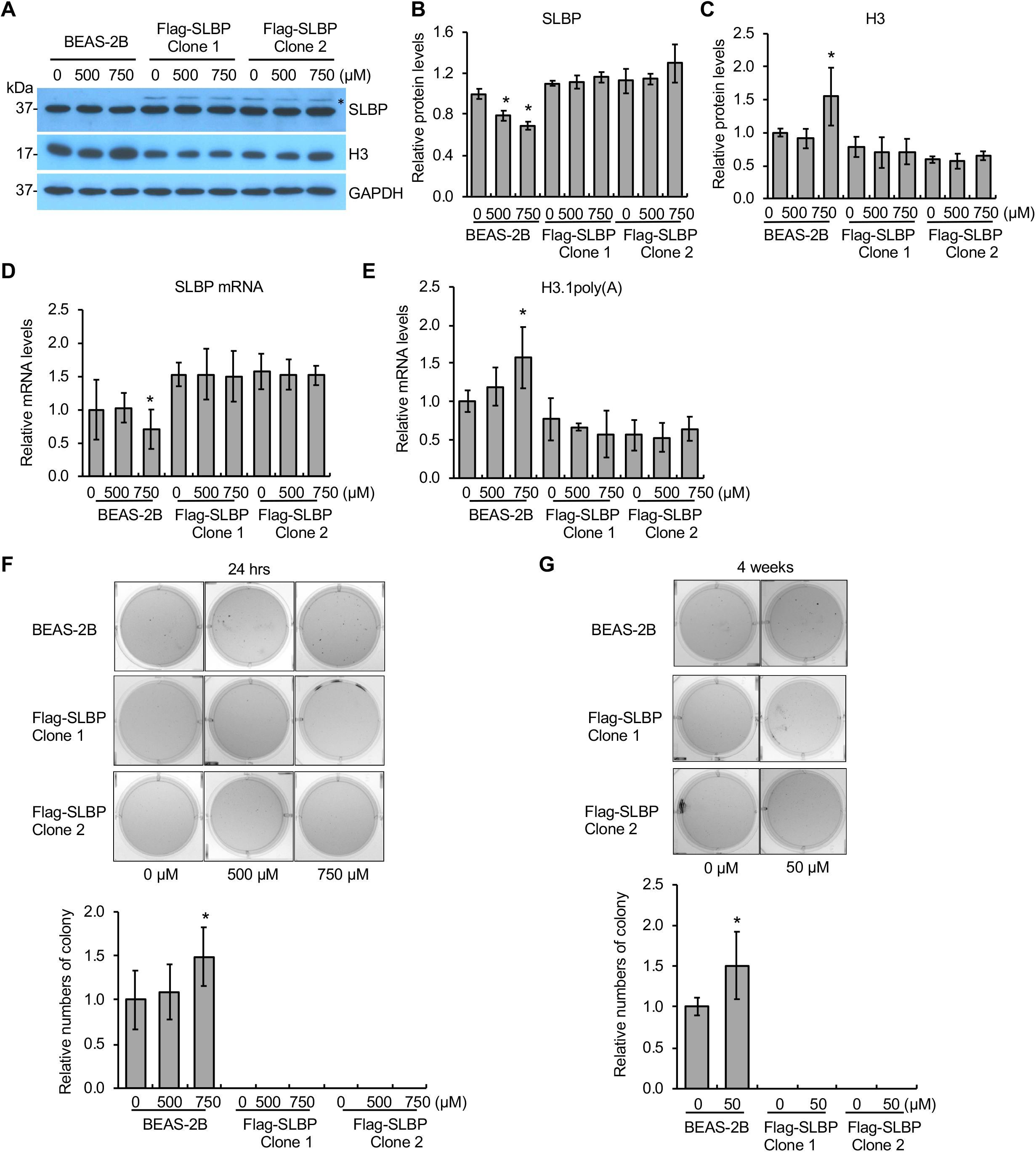
Overexpression of SLBP rescues nicotine-induced cell transformation. **(A-C)** Overexpression of SLBP rescues downregulation of SLBP by nicotine exposure to 500 μM or 750 μM for 24 hrs. BEAS-2B cells as well as two SLBP-overexpressing clones that have been stably transfected with FLAG-tagged SLBP-expressing vector were treated with or without nicotine and then subjected to Western blot (A). The asterisk next to “SLBP” indicates the ectopic SLBP. The band intensities were quantified using ImageJ software and presented as bar graphs to show relative quantifications of SLBP (B) and H3 (C). GAPDH was used as an internal control. The controls in lane 1 were used as references. The data shown are the mean ± S.D. (n = 3). \**p* < 0.05 vs. control group. **(D and E)** RT-qPCR detects changes in *SLBP* mRNA level and polyadenylation of H3.1 mRNA following nicotine exposure to 500 μM or 750 μM for 24 hrs in BEAS-2B as well as SLBP-overexpressing cells, i.e., Flag-SLBP clone 1 and clone 2. mRNA levels for *SLBP* and polyadenylated H3.1 were normalized to *GAPDH*. The data shown are the mean ± S.D. (n = 3). \**p* < 0.05 vs. control group. **(F and G)** Soft agar assays. BEAS-2B cells as well as SLBP-overexpressing cells were treated either with 0, 500, and 750 μM nicotine for 24 hrs, or with 0 and 50 μM nicotine for 4 weeks, and then plated in soft agar, cultured for 6 weeks. The data shown are the mean ± S.D. (n = 3). \**p* < 0.05 vs. control group.

### Exposure to nicotine aerosols generated from e-cigs downregulates SLBP in normal human bronchial epithelial (NHBE) primary cells and in mice

We used liquid nicotine to demonstrate downregulation of SLBP and subsequent increase in polyadenylated histone H3.1 mRNA and H3 protein level in BEAS-2B cells. It is important to confirm the results with nicotine aerosols generated by e-cigs. The nicotine concentration of e-cig varies between 3 and 36 mg/ml with recent generations of e-cigs containing up to 60 mg/ml of nicotine (30). Thus, we generated e-cig aerosols from unflavored e-liquid, which contains 0 or 18 mg/mL nicotine with 50% PG and 50% VG (50:50 PG/VG). Filtered air (FA) was used as the control. Exposure of BEAS-2B cells to e-cig aerosols generated from e-liquid with 18 mg/mL nicotine, but not with 0 mg/mL nicotine, dramatically reduced the protein and mRNA levels for SLBP as compared to the FA control group (Fig. 9A, B and D). Exposure to e-cig aerosols with 18 mg/mL nicotine significantly increased the protein level of H3 (Fig. 9A and C) and polyadenylated H3.1 mRNA as compared to the FA control (Fig. 9E). By contrast, exposure to e-cig aerosols without nicotine had no effect on the levels of polyadenylated H3.1 mRNA and H3 protein (Fig. 9A, C and E). These data indicate that the loss of SLBP and gain of polyadenylated H3.1 mRNA can be induced not only by exposure to liquid nicotine, but also by nicotine aerosols generated by heating e-cig containing nicotine.

**Figure 9.**
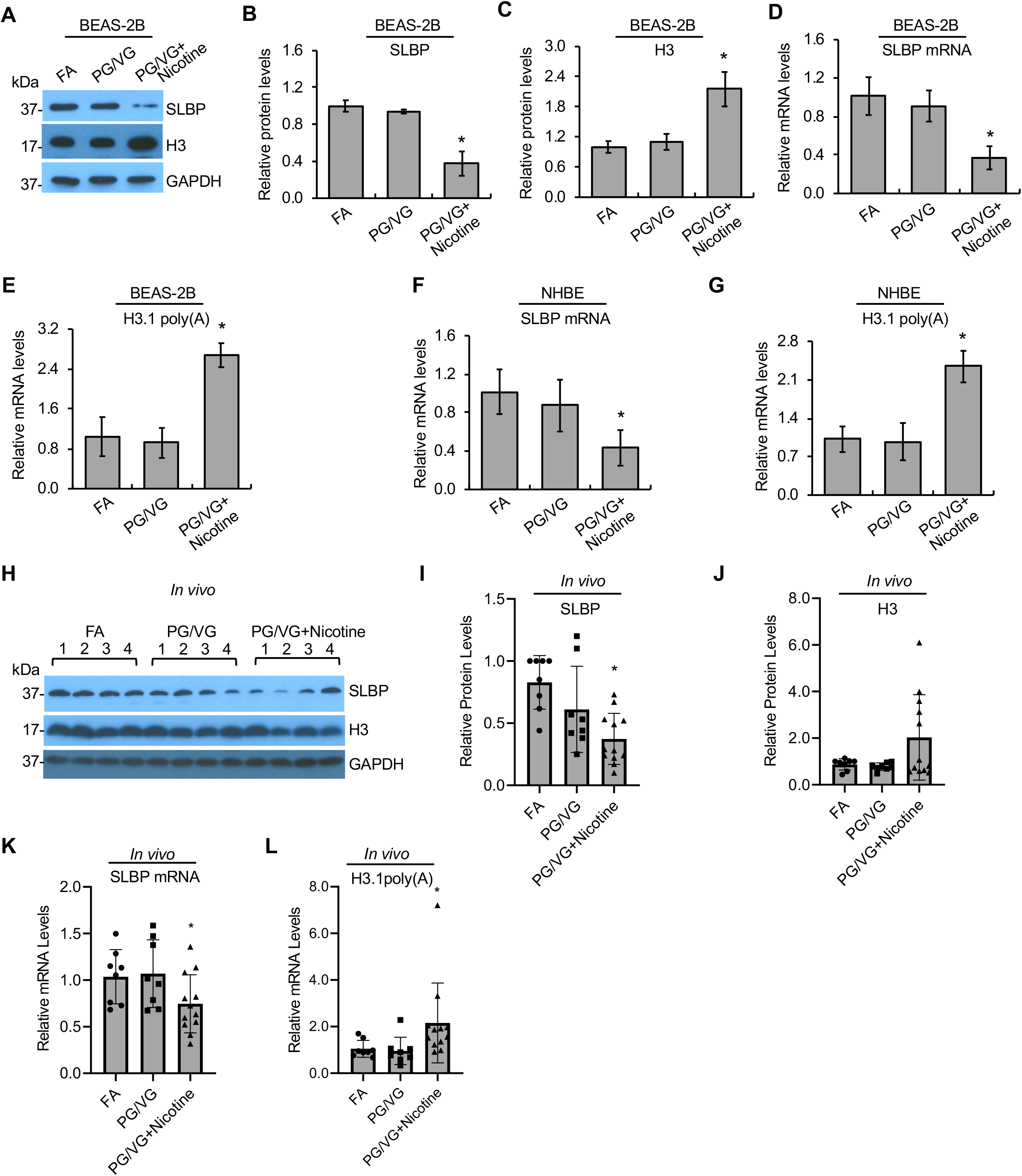
Downregulation of SLBP and polyadenylation of H3.1 mRNA by e-cig aerosols with nicotine in human lung primary cells and in mice. **(A-C)** Western blot detects changes in SLBP and H3 protein levels induced by e-cig aerosols in BEAS-2B cells. BEAS-2B ells were exposed to filtered clean air (FA), e-cig aerosols containing PG/VG and 0 mg/ml nicotine (PG/VG), or e-cig aerosols containing PG/VG and 18 mg/ml nicotine (PG/VG+Nicotine) followed by western blot analysis. The band intensities were quantified using ImageJ software and presented as bar graphs to show relative quantifications of SLBP (B) and total H3 (C). GAPDH was used as an internal control. The FA control group were used as references. The data shown are the mean ± S.D. (n = 3). \**p* < 0.05 vs. FA group. **(D and E)** RT-qPCR detects changes in SLBP mRNA and polyadenylated H3.1 mRNA levels induced by e-cig aerosols in BEAS-2B cells. The amount of polyadenylated H3.1 mRNA was measured by RT-qPCR using cDNAs synthesized with oligo (dT) primers. mRNA levels for SLBP and H3.1 were normalized to *GAPDH*. The data shown are the mean ± S.D. (n = 3). \**p* < 0.05 vs. FA group. **(F and G)** Changes in SLBP mRNA and polyadenylated H3.1 mRNA levels induced by e-cig aerosols in NHBE primary cells. mRNA levels for SLBP and polyadenylated H3.1 were normalized to *GAPDH*. The data shown are the mean ± S.D. (n = 3). \**p* < 0.05 vs. FA group. **(H-L)** Western blot detects changes in SLBP and H3 levels in lung tissues of mice exposed to e-cig aerosols. A/J female mice were exposed FA, PG/VG, or PG/VG plus 36 mg/ml of nicotine 5 days a week for 3 months. Lung tissues were collected and protein and mRNA levels were analyzed by Western blot (H-J) or RT-qPCR (K,L), respectively. GAPDH was used as an internal control. The data are shown for each individual mouse. Relative protein levels were calculated based on band intensity. Error bars represent S.D. (n=8 for FA and PG/VG groups; n=12 for nicotine exposure group). \**p* < 0.05 vs. FA group.

Next, we used normal human bronchial epithelial cells (NHBE) to validate our data in primary human lung cells. Since the cells grow very slowly with limited passage numbers, we were only able to collect samples for RT-qPCR assays. The level of *SLBP* mRNA was reduced by exposure of NHBE cells to e-cig aerosols with nicotine, but not by e-cig aerosols without nicotine as compared to the FA control (Fig. 9F). Furthermore, exposure of the primary human lung cells to nicotine aerosols increased the level of polyadenylated H3.1 mRNA by more than 2-fold (Fig. 9G). These data confirm the impact of nicotine on SLBP expression and polyadenylation of H3.1 mRNA in human lung primary epithelial cells.

Changes in SLBP expression and polyadenylation of H3.1 mRNA were also measured in lung tissues collected from female A/J mice exposed to FA, e-cigs without nicotine (PG/VG group) or e-cigs with 36 mg/mL of nicotine (PG/VG plus nicotine group). The mice were exposed 5 days a week for 3 months and analyzed 2.5 months later. Western blot and RT-qPCR results showed that both protein and mRNA levels of SLBP were decreased in the nicotine exposure group, but not in the PG/VG group, with statistical significance (*p* < 0.05) as compared to the FA control group (Fig. 9H, I and K). Moreover, the level of polyadenylated H3.1 mRNA was also increased in lung tissues of mice exposed to e-cig aerosols with nicotine (nicotine group), but not to e-cig without nicotine (PG/VG group) (Fig. 9L). The total H3 protein levels appeared to be increased in the nicotine group, but not in the PG/VG group as compared to the FA control, however, this did not reach statistical difference (Fig. 9H and J). These results suggest that nicotine exposure downregulates SLBP levels and induces polyadenylation of canonical histone mRNA *in vivo*.

## Discussion

In the present study, using immortalized human bronchial epithelial BEAS-2B cells, human primary NHBE cells, and an *in vivo* exposure, we demonstrate that nicotine exposure induces the loss of SLBP and the gain of polyadenylated canonical histone H3.1 mRNA. Treatment of BEAS-2B cells with liquid nicotine resulted in a decrease in both protein and mRNA levels of SLBP and the increase in the levels of H3.1 mRNA with poly(A) tail. Importantly, the same results were obtained with e-cig aerosols, which were generated by heating e-cig liquid containing nicotine. The results were not likely due to other components in the e-cig liquid, such as PG/VG, since the aerosols from e-cig without nicotine did not change the levels of SLBP and polyadenylated H3.1 mRNA as compared to the FA (filtered air) control. Moreover, the results were not likely BEAS-2B cells- and immortalized cells-specific, since exposure of the primary NHBE cells to nicotine aerosols also downregulated *SLBP* mRNA levels and increased the levels of polyadenylated H3.1 mRNA. These changes were seen in animal exposure studies as well. Mice exposed to the aerosols from e-cig with nicotine, but not from e-cig without nicotine, exhibited lower levels of SLBP protein and mRNA levels as well as higher levels of polyadenylated H3.1 mRNA in the lung tissues as compared to the FA control groups. Taking together, we conclude that nicotine exposure induces the SLBP depletion and polyadenylation of canonical H3.1 mRNA.

The loss of SLBP is associated with genomic instability. For example, the *slbp* mutants that express reduced level of SLBP in *Drosophila* exhibited increased frequency of DNA double-strand breaks and tetraploid karyotypes, as well as the frequency of loss of heterozygosity (LOH). The *slbp* mutants also facilitated the formation of heterochromatin (31). Moreover, knockdown of *slbp* in *C. elegans* caused defective chromosome condensation and segregation in embryos (32). While SLBP is involved in the most processes of biosynthesis of canonical histones, including their pre-mRNA processing, mRNA stability, nuclear export, and translation. Aberrant 3’ processing, i.e., polyadenylation of canonical histone mRNAs appeared to be the major mechanism for SLBP-associated genomic instability, since overexpression of polyadenylated H3.1 mRNA led to deregulation of cancer-associated genes, including lung-cancer related genes, aberrant cell cycle progress as well as chromosome aneuploidy and aberrations (17). Not surprisingly, both knockdown of SLBP and polyadenylation of H3.1 mRNA were able to enhance anchorage-independent growth of BEAS-2B cells, indicating a critical role for the loss of SLBP and subsequent acquisition of a poly(A) tail at the 3’ end of H3.1 mRNA in cell transformation (17). This current study showed that while nicotine exposure decreased the level of SLBP, overexpression of SLBP was able to prevent the nicotine-induced cell transformation. This effect was likely attributable to reduced induction of polyadenylated H3.1 mRNA, since overexpression of SLBP attenuated nicotine-induced polyadenylation of canonical histone mRNAs. We propose that the loss of SLBP and the gain of polyadenylated canonical histone mRNAs, in particular H3.1 mRNA, may represent a novel mechanism for nicotine toxicity and cell transformation.

Nicotine acts primarily by activating nAChRs, which are expressed not only in neuronal cells, but also in non-neuronal epithelial and endothelial cells (7,33). Deregulation of the nAChRs is often observed in many cancers, including lung cancer. For example, the upregulation of α7-nAChR has been shown to activate different downstream pathways, thereby stimulating the proliferation and migration of cancer cells in lung tissue (25,26). In our study, immunofluorescence staining and western blot analysis revealed upregulation of α7-nAChR following nicotine treatment. α7-nAChR was required for nicotine-induced SLBP downregulation since an inhibitor of α7-nAChR, but not of α3/α4-nAChRs, attenuated nicotine-induced loss of SLBP. This was further supported by the observation that the SLBP level was not changed by nicotine exposure in the α*7-nAChR* knockdown cells by siRNA. Our study further demonstrated that activation of α7-nAChR and subsequent depletion of SLBP are required for nicotine-mediated cell transformation. Tang et al. reported that nicotine causes lung carcinogenesis in mice through inducing DNA damage and inhibiting DNA repair (3). Our data suggested that non-genotoxic function of nicotine, i.e., activation of α7-nAChR and subsequent downregulation of SLBP and polyadenylation of canonic histone mRNAs, might be another cause or significant contributor to the nicotine-induced carcinogenesis. Another implication of this study is that even without diffusion to the cells, nicotine may play an important role in nicotine-induced toxicity/carcinogenicity *via* binding to and activating certain receptors on cell membrane. Interplays between nicotine-induced activation of nAChR and DNA damage as well as inhibition of DNA repair need to be further investigated in the future.

Previous studies have demonstrated that activation of nAChRs can trigger a varieties of downstream signal pathways, including PI3K/AKT pathway. The PI3K/AKT pathway is the most commonly altered signaling pathway in human cancers (34). Our data showed that nicotine induced phosphorylation of AKT at S473 but not at T308. This was different from the previous results by West et al. where they showed that both S473 and T308 of AKT were phosphorylated by nicotine treatment (35). Differences in the cell types and thus the composition of nAChR subunits, and doses and durations of nicotine exposures were likely attributable to the observed differences between two studies. In fact, in our study, the phosphorylation of AKT was induced by nicotine *via* α7-nAChR in BEAS-2B cells, whereas in their study α3/α4-nAChRs were shown to be involved in the nicotine-induced rapid AKT phosphorylation in normal human epithelial cells (NHBEs) and small airway epithelial cells (SAECs) (35). Moreover, the phosphorylation of AKT at T308 appeared to be a transient change with peak levels at 15-30 mins post treatment (35), which might be another reason why we did not see the change as we measured the phosphorylation status of AKT after 24 hrs of treatment. Our data indicate that nicotine-induced activation of PI3K/AKT pathway might be required for downregulation of SLBP, since SLBP level was reversed by pretreating the BEAS-2B cells with LY294002, a PI3K inhibitor. While mechanisms that underlie PI3K/AKT-induced SLBP depletion need further investigation, it is possible that AKT may downregulate the SLBP level both by transcriptional deregulation and by proteasome-mediated protein degradation process given that AKT can target numerous functional protein classes including transcription factors and E3-ubiquitin ligases among many others.

In normal cells, the SLBP levels are mainly regulated by a posttranslational mechanism. Phosphorylation of SLBP at T61 by cyclin A/CDK1 and T60 by CK2 are critical for degradation of SLBP by the SCF (SKP1-CUL1-F-box protein) ubiquitin ligase complex (36). Our siRNA knockdown experiments demonstrated that the SLBP loss induced by nicotine was completely reversed by CDK1 knockdown and also by CDK2 knockdown, but to a lesser extent, indicating that both CDK1 and CDK2 are required for nicotine-induced SLBP downregulation. Interestingly, nicotine-induced p-AKT^S473^ was not changed by CDK1 knockdown but was prevented by CDK2 knockdown. These results indicate that CDK1 might downregulate SLBP expression AKT-independently probably through direct phosphorylation of SLBP, while CDK2 might be involved in SLBP downregulation through activating AKT. It is known that CDK2/cyclin A2 phosphorylates AKT at both S477 and T479, triggering the canonical AKT-pS473 phosphorylation to promote AKT activation in response to several upstream signals (37). It would be interesting to test if nicotine exposure induces phosphorylation of AKT at S477 and T479 through CDK2, which is required for nicotine-induced SLBP depletion.

It has been known that the phosphorylation of SLBP by the tumorigenic potential protein kinase CK2 is primed by the phosphorylation by cyclin A/CDK1, both are required for SLBP degradation (38). We demonstrate here using an inhibitor that nicotine-induced downregulation of SLBP requires activation of CK2. This was not surprising given that it was well known that CK2 can directly phosphorylate SLBP, inducing proteasome-mediated SLBP degradation. Using siRNA knockdown experiments, we further showed that CK2α1, but not CK2α2, is necessary for the decreased SLBP expression following nicotine exposure. It has been shown that CK2 can specifically activate AKT at Ser129 (S129), thereby increasing association of AKT with Hsp90, which has the protective effect on T308 phosphorylation of AKT (39,40). The CK2 catalytic α2 subunit can be more important than the α1 subunit for AKT S129 phosphorylation and AKT downstream signaling (41). Given that in our studies nicotine-induced phosphorylation of AKT at S473 and that the phosphorylated of AKT at S473 was attenuated by the CK2α1 knockdown rather than the CK2α2 knockdown, CK2 is unlikely activating PI3K/AKT pathway through inducing phosphorylation at S129. It is known that CK2 can indirectly regulate AKT activity. For example, CK2 inhibits PTEN phosphatase activity, causing AKT activity; CK2 phosphorylation of mTOR kinases enhanced the AKT upstream activator mTORC2, which is responsible for S473 phosphorylation of AKT (42). Thus, in addition to direct phosphorylation of SLBP by CK2 and subsequent degradation, indirect activation of AKT pathway by CK2 might also be involved in the nicotine-induced SLBP depletion.

Upregulation of CDK1/CDK2 and CK2 by nicotine was dependent on α7-nAChR. α7-nAChR was also required for nicotine-induced activation of PI3K/AKT pathway. Our findings lend support to the importance of nAChR in nicotine toxicity and potential carcinogenicity. It implies that nicotine may change cell functions and cell fate without entering the cells. Indeed, both inhibition of α7-nAChR by an inhibitor or by siRNA knockdown reversed nicotine-induced SLBP depletion and cell transformation. Since PI3K/AKT pathway and kinases like CDK1/2 and CK2 possess many downstream effectors, it is likely that nicotine may exert its effects not only through targeting SLBP but also other effectors. Nonetheless, the loss of SLBP and subsequent gain of polyadenylation of canonical histone mRNAs appeared to play an important role in nicotine-mediated cell transformation, as nicotine-induced anchorage-independent cell growth was rescued by overexpression of SLBP. We propose that nicotine causes SLBP depletion and polyadenylation of canonical histone mRNAs through activation of α7-nAChR and a series of downstream signal transduction pathways, involving PI3K/AKT, CDK1/2, and CK2. This mechanism may be critical for nicotine-induced bronchial epithelial cell transformation and carcinogenesis.

## Acknowledgements

We thank John Adragna and Terry Gorden for their assistance in cell exposure to e-cig aerosols. This work was partially supported by grants from the US National Institutes of Health: R01ES029359 and R01ES030583 (C.J.).

## Author Contributions

C.J conceived the study; Q.S. and C.J. designed the experiments; Q.S. and D.C. performed most of the experiments and data analysis; A.R. conducted animal exposure under supervision of J.Z and G.G; G.G, Q.S., and D.C. collected mouse lung tissues; Q.S. and C.J. wrote the manuscript with input from other co-authors.

