## Supplemental Figures for "Downregulation of Stem-loop binding protein by nicotine *via* α7-nicotinic acetylcholine receptor and its role in nicotine-induced cell transformation"

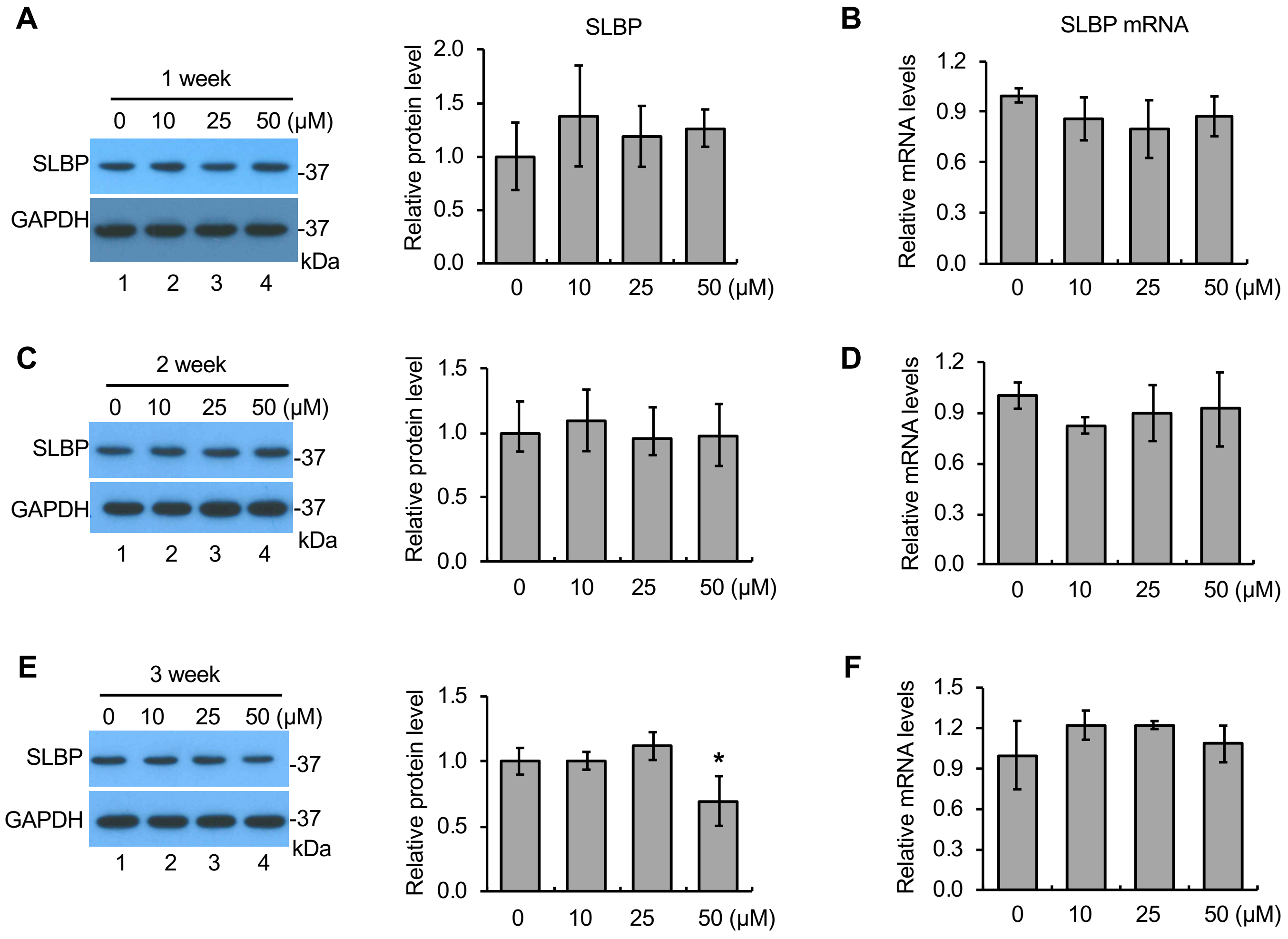


**Supplementary Figure 1. Changes in SLBP protein and mRNA levels in BEAS-2B cells following low-dose nicotine treatment for 1-3 weeks**

The SLBP protein and mRNA levels were measured by Western blot (A, C, and E) and RT-qPCR (B, D, and F), respectively, in BEAS-2B cells treated with (10 μM, 25 μM, and 50 μM) or without nicotine for 1- (A and B), 2- (C and D), and 3-weeks (E and F). GAPDH was used as an internal control. The band intensities (left panels in A, C, and E) were quantified and presented as bar graphs (right panels in A, C, and E). The controls in lane 1 were used as references. The data shown are the mean ± S.D. (n = 3). *$p < 0.05$ vs. control group.

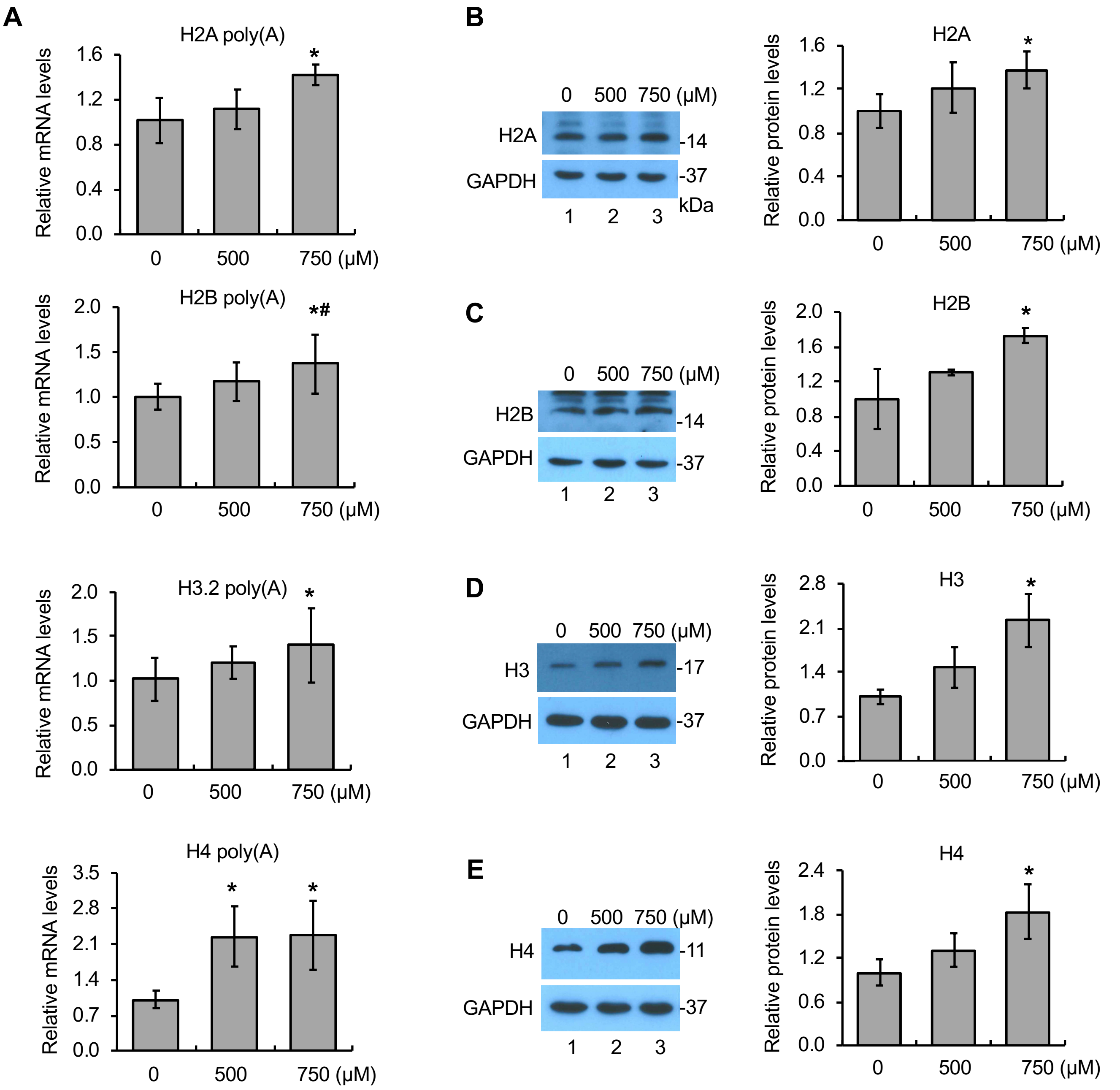


**Supplementary Figure 2. Polyadenylation of canonical histone mRNAs by nicotine**

The level of polyadenylated mRNAs for canonical histones H2A, H2B, H3.2, and H4 (A) as well as proteins for H2A, H2B, H3, and H4 (B-E) were determined by RT-qPCR and Western blot, respectively, in BEAS-2B cells treated with (500 μM and 750 μM) or without nicotine for 24 hrs. GAPDH was used as an internal control. The band intensities (left panels in B-E) were quantified and presented as bar graphs (right panels in B-E). The controls in lane 1 were used as references. The data shown are the mean ± S.D. (n = 3). *$p < 0.05$ vs. control group.

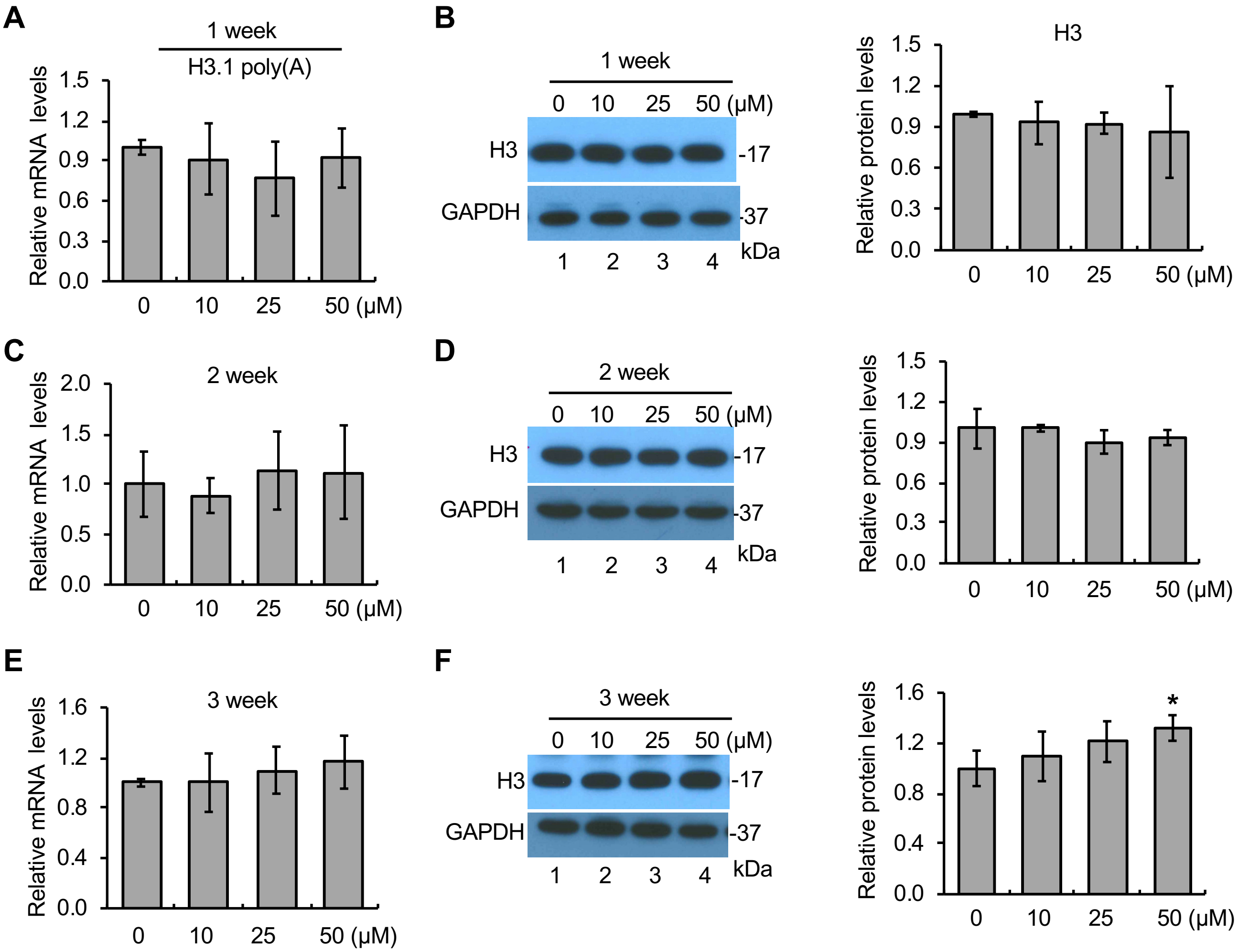


**Supplementary Figure 3. Polyadenylation of canonical histone H3.1 mRNA by low-dose nicotine treatment for 1-3 weeks**

The level of polyadenylated H3.1 mRNA (A, C, and E) and total H3 protein level (B, D, and F) were determined by RT-qPCR and Western blot, respectively, in BEAS-2B cells treated with (10 μM, 25 μM, and 50 μM) or without nicotine for 1- (A and B), 2- (C and D), or 3-weeks (E and F). GAPDH was used as an internal control. The band intensities (left panels in B, D, and F) were quantified and presented as bar graphs (right panels in B, D, and F). The controls in lane 1 were used as references. The data shown are the mean ± S.D. (n = 3). *$p < 0.05$ vs. control group.

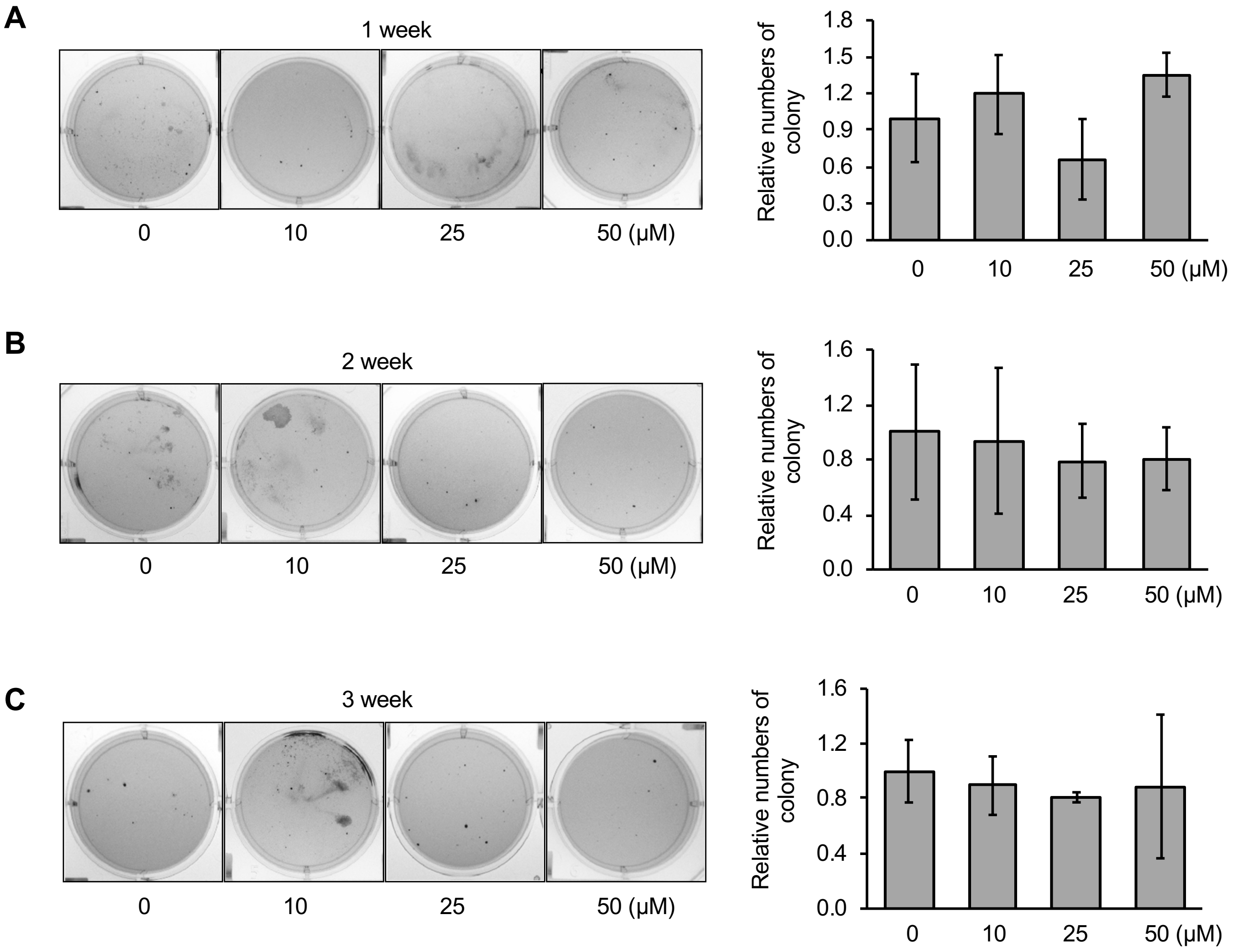


**Supplementary Figure 4. Anchorage-independent cell growth induced by low-dose nicotine treatment for 1-3 weeks**

BEAS-2B cells treated with (10 μM, 25 μM, and 50 μM) or without nicotine for 1- (A), 2- (B), or 3-weeks (C) were subjected to soft-agar assays. The cells were plated in soft agar and cultured for 6 weeks. The data shown are the mean ± S.D. (n = 3). $^{*}p < 0.05$. vs. control group.

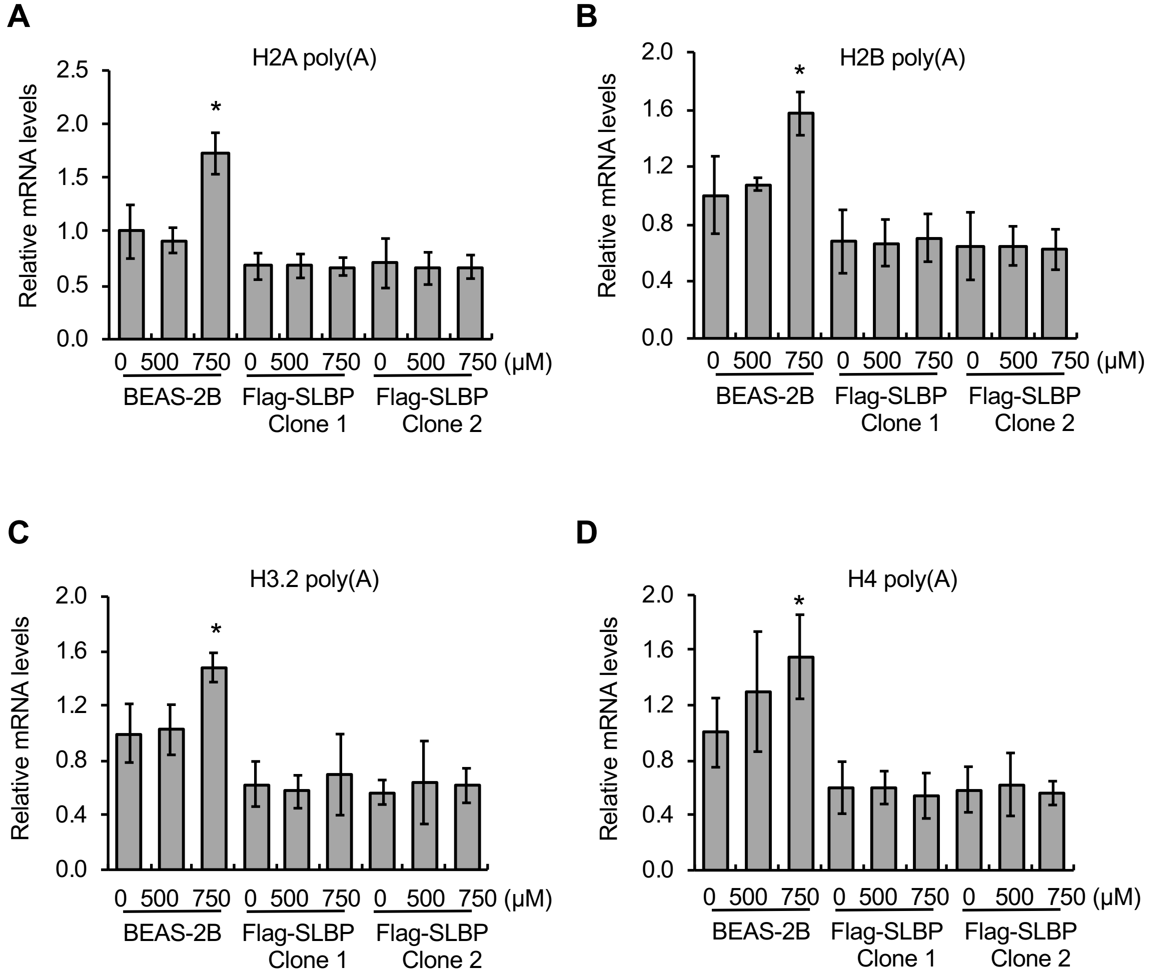


**Supplementary Figure 5. Overexpression of SLBP rescues polyadenylation of canonical histone mRNAs for H2A, H2B, H3.2, and H4**

RT-qPCR was used to determine the levels of polyadenylated mRNAs for H2A (A), H2B (B), H3.2 (C), and H4 (D) in BEAS-2B cells and SLBP-overexpressing stable cell lines, i.e., Flag-SLBP clone 1 and clone 2. GAPDH was used as an internal control. The data shown are the mean ± S.D. (n = 3). *$p < 0.05$ vs. control group.
